# Experimental Removal of Niche Construction Alters the Pace and Mechanisms of Resistance Evolution

**DOI:** 10.1101/2025.03.27.645637

**Authors:** Lai Ka Lo, Nora K. E. Schulz, Helle Jensen, Joachim Kurtz

## Abstract

Niche construction, a central eco-evolutionary process in which organisms modify their environment, is hypothesised to enhance evolutionary adaptability through feedback between genetic inheritance and the lasting effects of environmental change—known as ecological inheritance. However, while theoretically intriguing, direct empirical support for the effect of niche construction on adaptation is lacking. Red flour beetles modify their environment by releasing quinone-rich stink gland secretions, a form of external immunity. Here, we test in an evolution experiment how experimental removal of niche construction—using RNAi of a key gene needed to produce stink gland secretions—affects the beetles’ adaptation to the bacterial entomopathogen *Bacillus thuringiensis.* In each generation, beetles inhabited conditioned flour either with regular or drastically reduced stink gland secretion content. Compared to the non-exposed control, all bacteria-exposed regimes rapidly evolved bacterial resistance within only six generations. However, beetles evolving with intact niche construction acquired resistance as soon as after three generations, indicating that adaptation was initially facilitated by niche construction. RNAseq of evolved beetles showed that gene expression differed strongly between the selection regimes, revealing that the mechanisms underlying resistance were dependent on niche construction. Thus, our findings provide urgently needed empirical evidence on the role of niche construction for resistance evolution and its potential genetic basis.

## Main

Organisms often engage in niche construction by actively altering abiotic and biotic aspects of their environments to maintain or increase its suitability for themselves and their offspring ^1–3^. Prime examples include earthworms and plants modifying soil structure and chemistry, beavers and dung beetles building their nests, and termites cultivating beneficial fungi ^4–7^. A recent meta-analysis on the niche construction done by invertebrates across all continents, also demonstrated how hugely beneficial their ‘engineering work’ can be for entire ecosystems^8^. Niche construction theory proposes that organisms can shape their environment in ways that influence their and their offsprings’ selective pressure, challenging the long-standing view of evolution where organisms passively adapt to their environments via natural selection. As successive generations develop in and adapt to the altered environment, they further contribute to the niche construction through behaviours or physiology that enhance phenotype-environment fit^5,9,10^. Through ecological inheritance these changes accumulate and are passed on, alongside genes, influencing trait-fitness relationships under selection. Therefore, niche construction can drive the divergence of niches and local adaptations across and within populations and can promote the maintenance or emergence of phenotypic variation as a response to diverse and ever-changing environments ^11–15^.

Despite extensive discussion, researchers almost exclusively have relied on mathematical models^16–18^ and case studies of wild animals^19^ to investigate the impact of niche construction on evolution. Such lack of empirical support has raised criticism, such that the role of niche construction in evolution has remained controversial. More experimental work includes microbial populations evolving dependency on their modified chemical environment and creating new ecological niches^20^, or dung beetle mothers and larvae constructing their own developmental niches which can shape phenotypic variation and fitness outcomes^6^. However, as far as we are aware, the very critical empirical investigation on how niche construction affects selection is still lacking, especially in the context of host-parasite interactions, a key selective force for host survival and reproductive success. Besides their intrinsic immune system, hosts often reduce infection risk through niche construction (e.g., secreting antimicrobials to modulate surrounding microbiota) or niche choice (e.g., relocating to areas with fewer parasites)^21^. Host resistance is typically a polygenic trait shaped by genetic and environmental interactions, which lead to niche individualisation based on immune experience^22,23^. Thus, the fundamental question arises, how does niche construction modulate the host’s adaptation to a parasite and consequently its evolvability?

To experimentally study the effect of niche construction on resistance evolution and host adaptation to a pathogen, we performed an evolution experiment using the well-established host-parasite model of the red flour beetle, *Tribolium castaneum* and the entomopathogenic bacterium, *Bacillus thuringiensis tenebrionis* (*Btt*). *T. castaneum* adults condition their flour substrate with their antimicrobial stink gland secretions (SGS), a niche-constructing trait that can confer external immunity for themselves and their offspring^14,24,25^. These secretions regulate microbial niches, favouring the growth of yeast as a food source, while inhibiting other microbes^26,27^. Stink gland secretion is a heritable trait influenced by genetics^25,28,29^ and environmental factors such as parasite prevalence^29,30^, resource availability^31^ and the individual’s immune experiences^32^. The SGS-conditioned flour is passed on to conspecifics and offspring via ecological inheritance, and it can have fitness consequences^25^. In our study, we experimentally manipulated this niche constructing trait via RNA interference^33^ and evolved the focal beetles in differentially constructed niches conditioned by adults with either functional or impaired SGS production, before and after pathogen selection. We sought to determine (i) whether niche construction affects host adaptation to pathogens, (ii) how fitness outcomes differ between selection populations evolving in different niches, and (iii) the associated transcriptomic changes underlying adaptive responses to *Btt* when evolving in different niches.

## Results

We experimentally disrupted stink gland secretion in *T. castaneum* by RNAi-mediated knockdown of the *Drak* gene^33^ (see Supplementary Methods and Results for details) and used flour conditioned by the treated or control beetles as differentially constructed niches (Figure 1). Knockdown beetles showed significantly reduced *Drak* expression for at least 28 days (Fig. S1), produced minimal SGS (Fig. S2) and displayed weaker antibacterial activity (Fig. S3). For experimental evolution, we placed untreated beetles into these differentially conditioned niches from oviposition onwards until adult emergence, only interrupted by five days of oral *Btt* exposure as selection pressure during the larval stage. To enable reciprocal eco-genetic feedback during experimental evolution, siblings of each parental generation served as ‘niche constructors’, conditioning the flour to provide co-evolving niches for the next generation.

**Figure 1.**
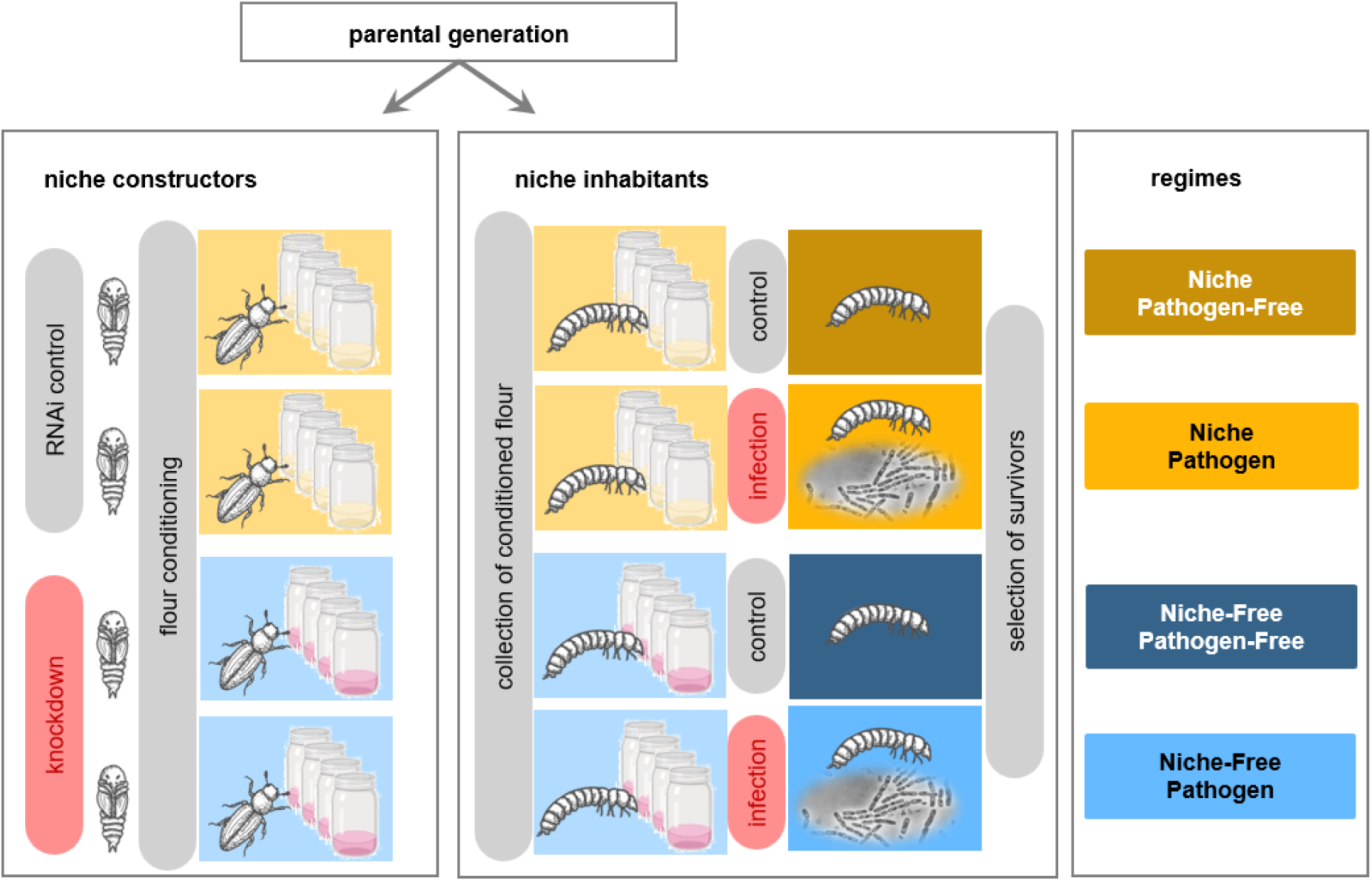
Scheme of evolution experiment with four selection regimes. Beetles raised in constructed niches (‘*niche’*; injection of dsRNA of the *Gfp* gene as RNAi control) or non-constructed niches (denoted ‘*niche-free’* for simplicity; injection of dsRNA of the *Drak* gene to achieve knockdown) were exposed to *Btt* spores (‘*pathogen’*) or not (‘*pathogen-free’*). Each selection regime consisted of four replicate populations with 100 individuals each. Niche constructors were siblings of parents of niche inhabitants and provided differentially conditioned flour for the two niche types.

### Constructed niches facilitate rapid evolution of resistance against *Btt*

To determine whether evolution in the constructed niche confers a post-infection survival advantage, we exposed F2 offspring of generations three, six and nine from all selection regimes to a bacterial challenge. Their survival rates were then compared with those of the *niche / pathogen-free* control selection regime.

Already after three generations of selection, the *niche / pathogen* regime showed significantly higher survival than the control *niche / pathogen-free* regime (Figure 2A, Hazard Ratio (HR) ±SE = 0.70 ±0.11, p < 0.001), while larvae from the other pathogen selected regime, *niche-free / pathogen* did not survive significantly better than the control (HR±SE = 0.85 ±0.10, p=0.12). There was also no difference in survival probabilities between the two regimes without *Btt* selection (HR±SE = 0.99 ±0.10, p=0.92). After six generations, both pathogen selected regimes showed increased survival compared to the control regardless of the presence of the constructed niche (*niche / pathogen*: HR±SE = 0.76 ±0.10 SE, p<0.01; *niche / pathogen-free*: HR±SE = 0.68 ±0.10, p < 0.001). Finally, at the time of the last phenotypic readout for generation nine, the hazard ratio for the probability of dying in the *niche / pathogen* regime went down to 0.39 (±0.13 SE, p<0.001) in relation to the *niche / pathogen free* control regime (Figure 2A). Also, the *niche-free / pathogen* regime had a similar survival probability (HR±SE = 0.50 ±0.12, p < 0.001), demonstrating that resistance can evolve independent of the constructed niche within the time frame of nine generations. Surprisingly, after nine generations of selection the *niche-free / pathogen-free* regime also showed improved survival of *Bt* infection despite never being exposed to the pathogen (HR±SE = 0.80 ±0.11, p < 0.05).

**Figure 2.**
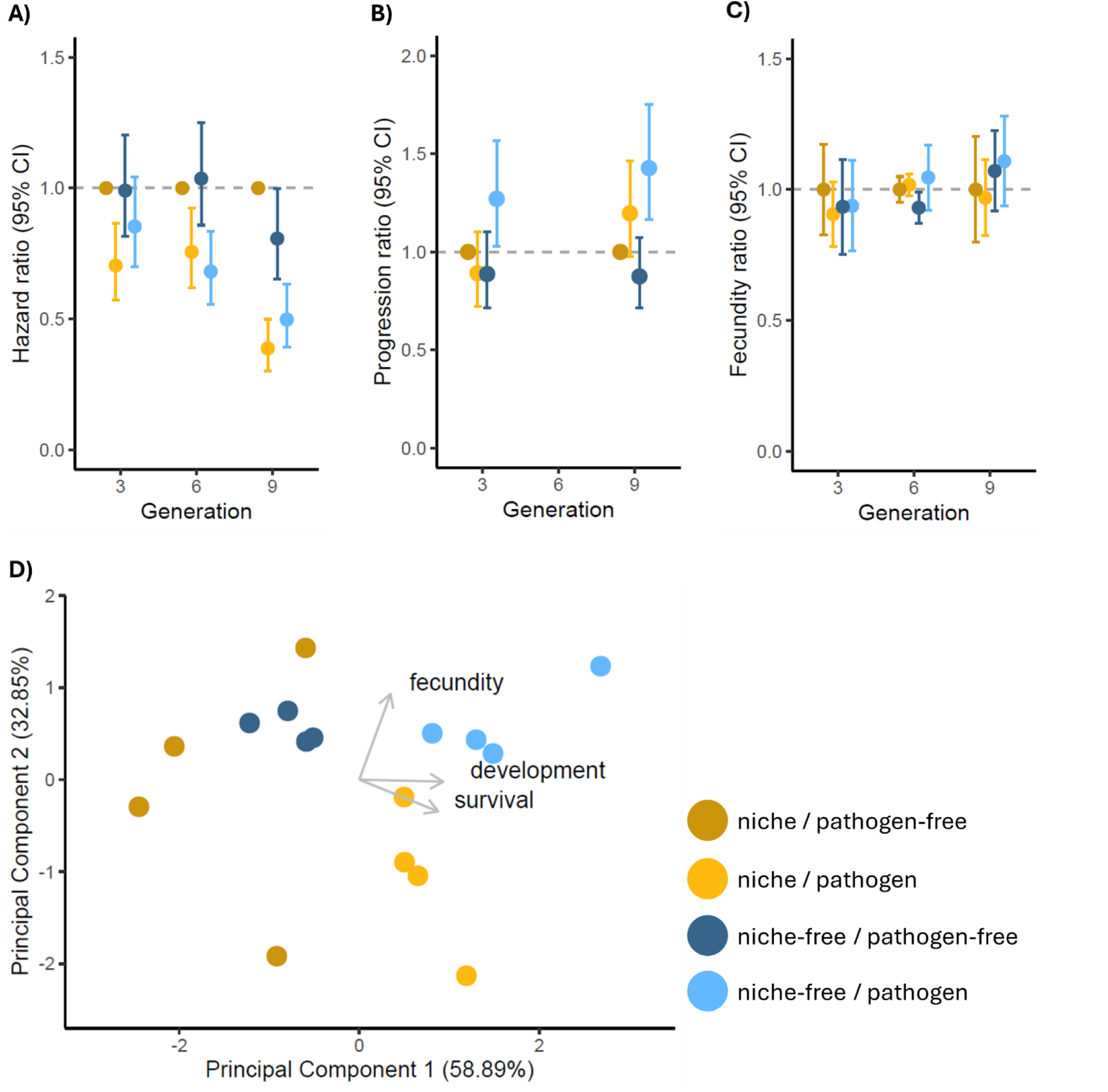
Resistance and life history traits of evolving populations. **A)** Hazard ratios of F2 offspring for dying from *Btt* exposure for all selection regimes compared to *niche / pathogen-free* regime, the control selection regime. Depicted is the median hazard ratio of the four replicate lines for the selection regimes with 95% confidence intervals. A hazard ratio smaller than 1 means that the treatment group experienced a lower mortality probability compared to the control group. Per line 96 larvae were exposed to *Btt* and another 96 served as control (data not shown) **B)** Developmental progression ratio showing the probability of F2 larvae to pupate between 19 to 32 days post-oviposition relative to the control regime with 95% confidence intervals. Ratios higher than 1 indicate faster development to pupal stage compared to the control. Given is the median for the four replicate lines per regime, with n = 48 individuals per replicate line. We do not have data for generation six. **C)** Early life fecundity of the F2 generation relative to the control regime. Shown is the mean number of offspring per pair, averaged across all four replicate lines with 95% confidence intervals (n = 20 pairs per replicate line). **D)** Biplot showing the results of a principal component analysis (PCA; variance explained by PC1 and 2) based on host life history traits, “Survival” (survival probability upon bacterial exposure), “Developmental time” (day to pupation) and “Fecundity” (number of live larvae produced) across selection lines in generation 9, with each dot representing one of the four replicate lines.

### *Btt* exposure in the absence of a constructed niche led to faster development

To test for potential effects of the evolution treatments on development, we measured the time to pupation of the F2 offspring for generations three and nine. In generation three, only the *niche-free / pathogen* regime led to significantly faster development than the control regime (Figure 2B, Progression ratio (PR±SE) =1.27 ±0.11, p < 0.05), suggesting that beetles accelerate development to escape the susceptible larval stage under the combined pressure from pathogen secretion and reduced niche construction. This difference increased in generation nine (PR±SE = 1.43 ±0.10, p < 0.001; Figure 2B).

### Little effects of selection regimes on fecundity

To investigate potential effects of the selection regime on reproductive fitness, we quantified the number of larvae two weeks after mating in the F2 offspring of the selection lines. We observed only weakly reduced offspring numbers by 7% (±1.8% SE, p<0.05) in the *niche-free/ pathogen-free* regime in generation six. Combining regimes according to niche type (*niche* or *niche-free*), revealed a minor, but significantly higher fecundity in the *niche-free* regimes (Figure 2c, χ^2^ = 5.889, df= 1, p = 0.015), while there was no effect of pathogen exposure (χ^2^ = 0.57, df= 1, p = 0.45) and no interaction between pathogen exposure and niche (χ^2^ = 1.267, df= 1, p = 0.26) on fecundity.

### Life history traits vary with niche and *Btt* selection regimes

To explore how populations from different regimes might differ in terms of their life history, we jointly analysed survival, developmental time and fecundity in a PCA for generation nine. Survival and development were captured by the first principal component (PC1), accounting for 58.89% of the variance (Figure 2D) whereas fecundity contributed largely to the second principal component (PC2) explaining 32.85 % of the variance (Figure 2D). PERMANOVA revealed that the presence or absence of the constructed niche explained a significant 40.8% of the variance (p < 0.01), whereas *Btt* selection accounted for only a non-significant 6.2% (p = 0.35). We also tested whether total amounts of SGS differed among the selection regimes, but did not find any strong differences (Fig. S4).

### Niche construction strongly impacts changes in gene expression in response to bacterial selection

To assess whether niche conditions led to divergent mechanisms of bacterial resistance, we compared gene expression in F2 larvae from generation nine across all regimes, both six hours post-*Btt* exposure (induced response) and without exposure (constitutive response). There were only three constitutive DEGs between the two *pathogen-free* regimes, demonstrating that the differences in niche construction alone did not lead to changes on the transcriptomic level (Fig. 3A). However, there were substantial differences in constitutive gene expression when the *Btt* selection was added to the picture. In both *pathogen* regimes a total of 288 genes are constitutively differentially expressed compared to the *pathogen-free* regimes, demonstrating considerable overlap in adaptation to the pathogen. However, there are also considerable differences in the response to the selection. In the presence of the niche only an additional 187 genes are differentially expressed, while this number more than triples with 623 DEGs in the *niche-free* regime. These over 800 DEGs in total showed a large overlap in gene ontology (GO) terms, with ‘peptidase activity’, ‘oxidation-reduction process’, and ‘proteolysis’ being most dominant (Figure 3B).

**Figure 3.**
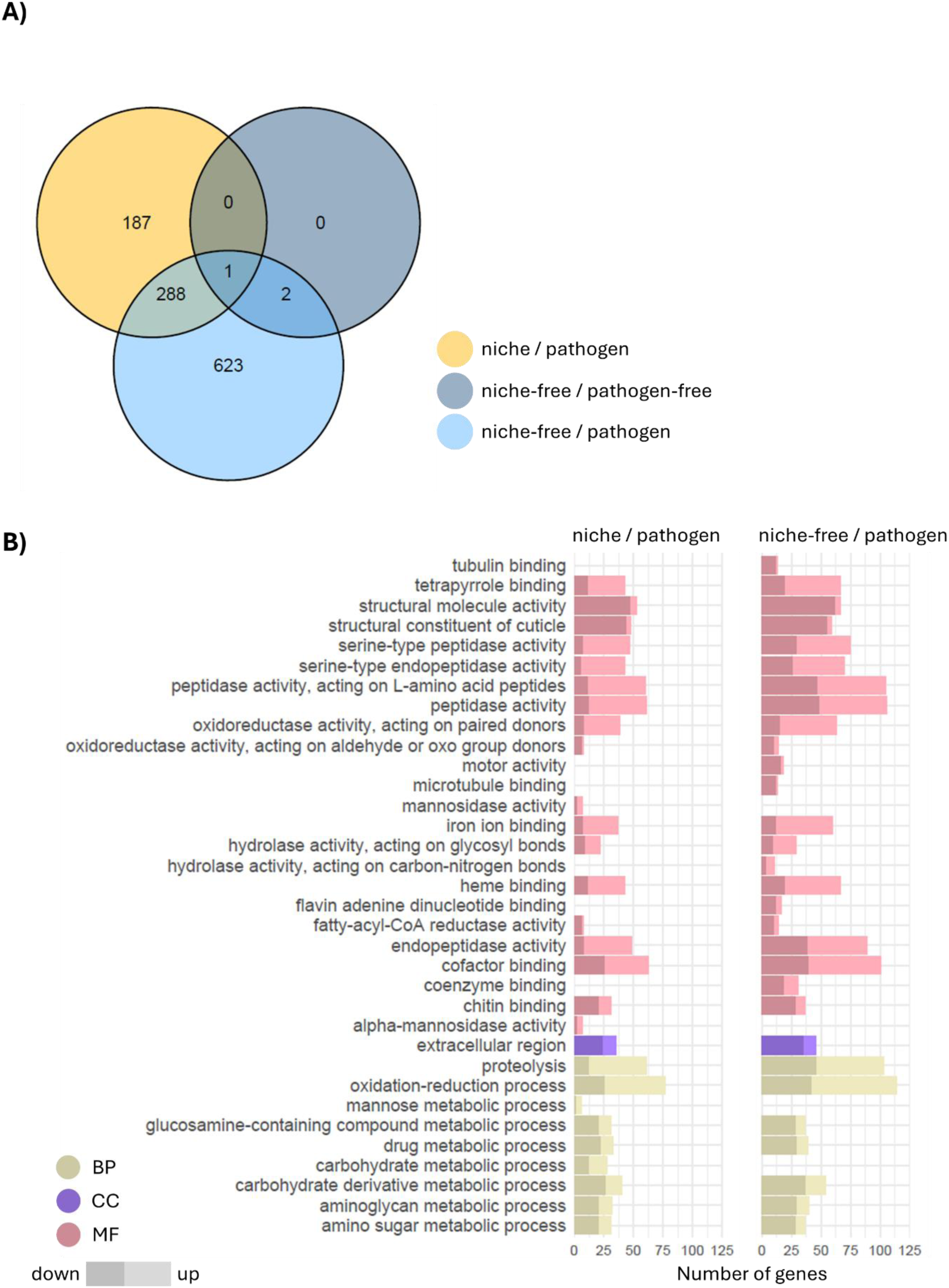
Differential gene expression between the regimes without *Btt* exposure. Results of the analysis with DEseq2 (padj<0.05, any Log-fold change). **A)** Venn diagram of DEGs for the three regimes relative to the control regime *niche / pathogen-free* without bacteria exposure. **B)** Enriched GO terms (padj<0.05) in *niche / pathogen* and *niche-free / pathogen* regime compared to *niche / pathogen free* control. Shown are up to ten of the top GO terms for each category and regime (BP= biological process, CC= cellular component, MF= molecular function, empty rows indicate non-significant results).

When we compare the induced responses between the regimes six hours after a bacterial exposure, the picture reverses to much stronger changes in expression in the *niche / pathogen* regime, where 562 genes are uniquely differentially expressed in this regime (Figure 4A). Again, there was considerable overlap in enriched GO terms, with shared prominent terms including ‘oxidation-reduction process’, ‘cofactor binding’, and ‘peptidase activity’ (Figure 4B). The majority of the DEGs in these GO terms were upregulated, hinting at a stronger and faster response to the spores in the *Btt* selected regimes, especially *niche / pathogen*. Lastly, there were only a handful of DEGs between the *niche-free / pathogen-free* and the control, which implies that the presence/absence of a constructed niche alone is not enough to cause evolved changes in gene expression profiles (Figure 4A).

**Figure 4.**
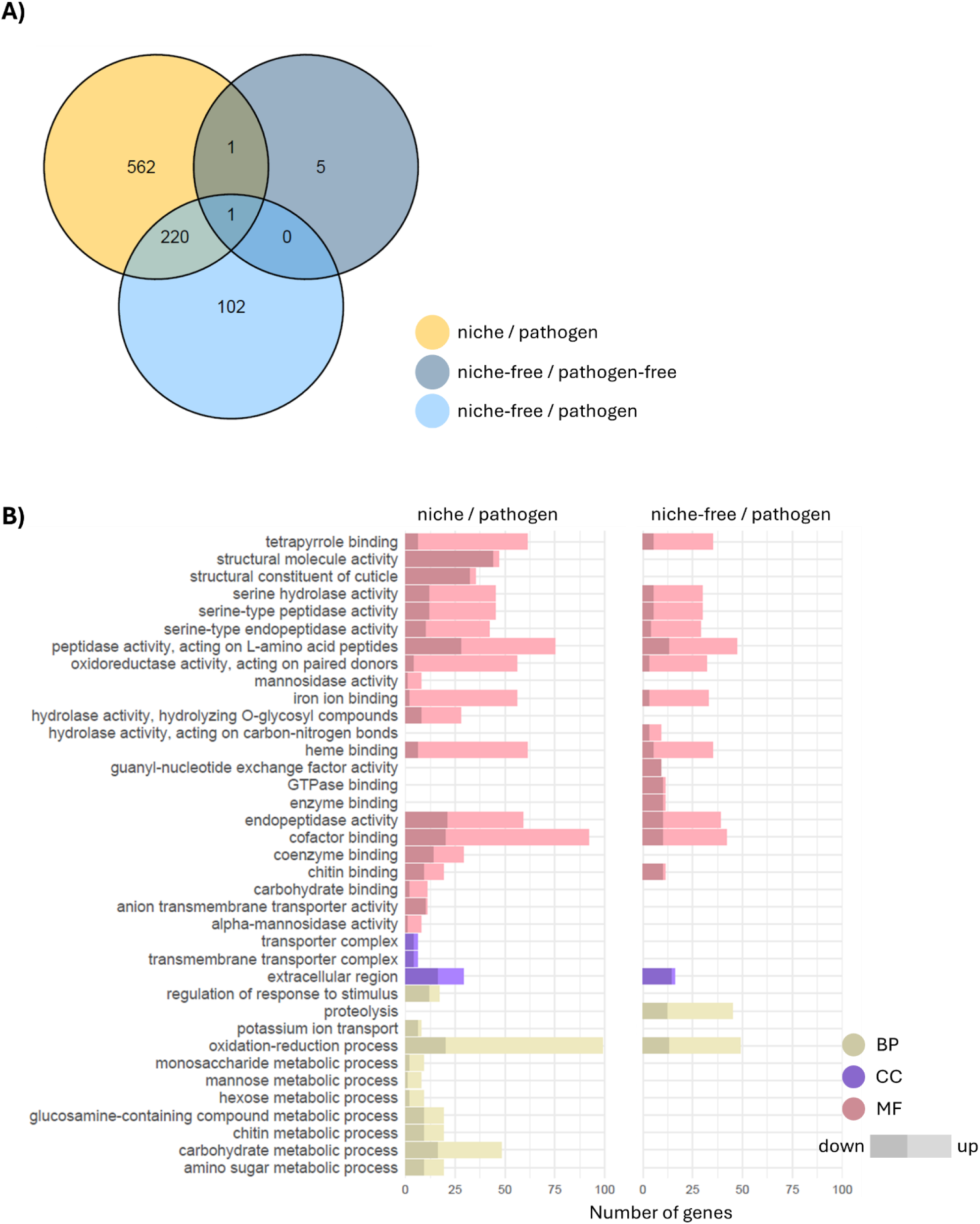
Differential gene expression between the regimes following *Btt* exposure. Results of the analysis with DEseq2 (padj<0.05, any Log-fold change). **A)** Venn diagram of DEGs for the three regimes relative to the control regime *niche / pathogen-free* six hours post bacteria exposure. **B)** Enriched GO terms (padj<0.05) in *niche / pathogen* and *niche-free / pathogen* regime compared to *niche / pathogen free* control. Shown are up to ten of the top GO terms for each category and regime (BP= biological process, CC= cellular component, MF= molecular function, empty rows indicate non-significant results).

### Minor differences in early response to *Btt* exposure between selection regimes

Next, we compared the differences in responses in gene expression to *Btt* exposure within selection regimes (Figure S5). The *niche-free / pathogen* regime showed the strongest response (346 DEGs), nearly double that of the next highest (*niche / pathogen*:182 DEGs). The DEG regulation showed no consistent directional trend.

GO term enrichment revealed broad overlap across regimes, with common categories including oxidation-reduction, proteolysis, and carbohydrate metabolism (Figure S6; Figure S7). Some unique enrichments — e.g., carboxylic acid metabolism in the niche/pathogen regime and glycolysis-related terms in the niche-free/pathogen-free regime — suggest minor metabolic differences. Peptidase and hydrolase activity were enriched across all regimes; “virion part” was enriched only in the niche regimes.

### Constitutive changes in gene expression vary by niche in the *Btt* selected lines

To further examine the variation in constitutive and infection-induced gene expression among the different evolved regimes, weighted gene co-expression network analysis (WGCNA) was performed separately between the non-exposed (Figure S7) and *Btt*-exposed (Figure S8) larvae from all four regimes. In samples without bacterial exposure, three modules (purple, magenta, and greenyellow) significantly negatively correlated with the absence of a constructed niches in the *pathogen* regime, while there were no significant correlations in the *pathogen-free* regimes (Figure 5). This suggests that genetic changes mediated by differently constructed niches were only apparent under constant bacterial selection.

**Figure 5.**
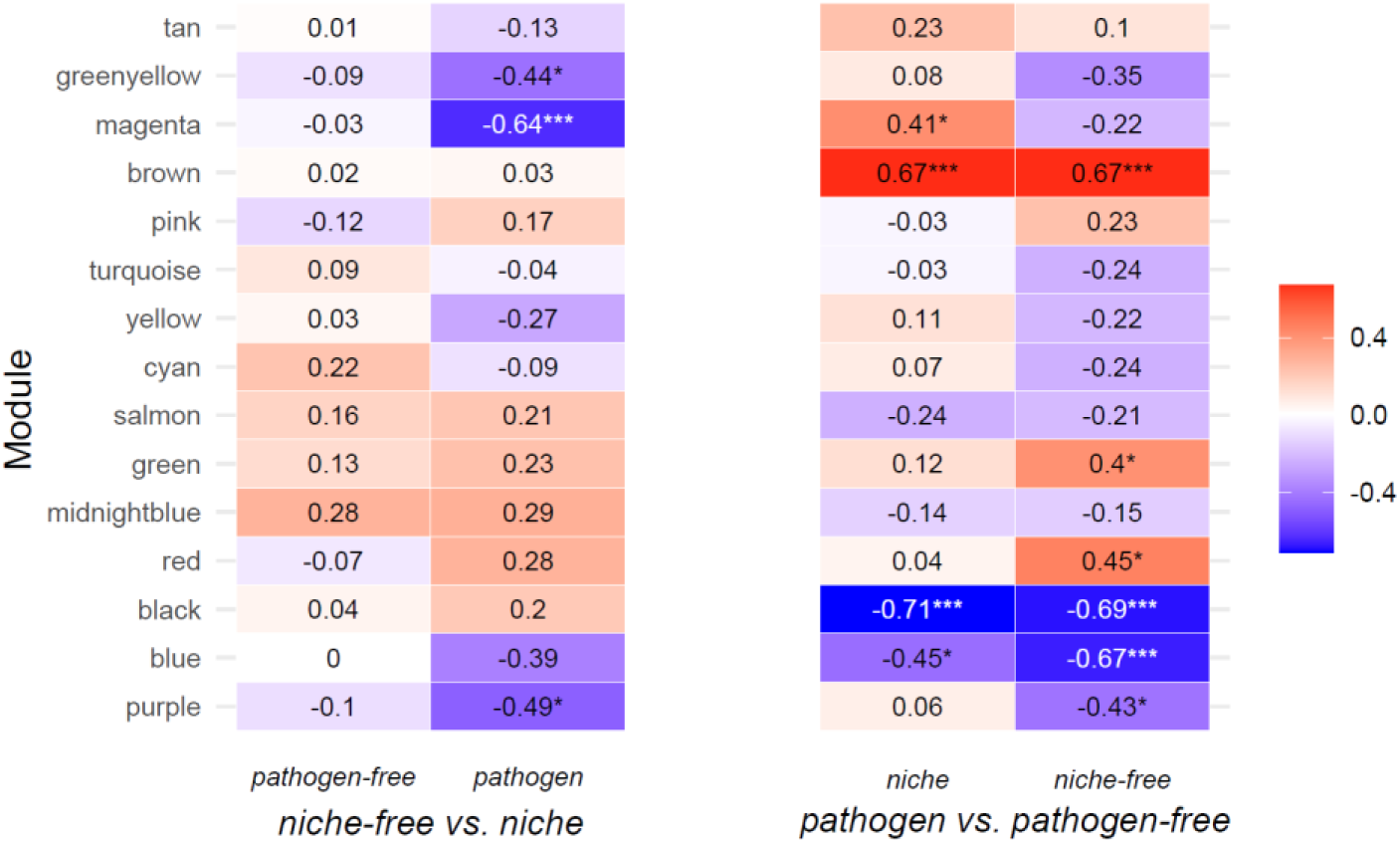
WGCNA for the control samples (without *Btt* exposure). Heat map of module-selection regime correlations. Cell colours indicate Pearson coefficients, and asterisks denote significance levels.

The magenta module is characterised by genes involved in endocytosis related to synaptic vesicle recycling, clathrin light chain binding, and biological adhesion (Figure 6A). The hub gene of this module is TC004152, “E3 ubiquitin-protein ligase HUWE1-like Protein” (HUWE1) which is crucial for DNA repair and mitigating replication stress^34^. Similarly, the purple module was significantly down-regulated in the *niche-free / pathogen* regime. This module includes genes vital for DNA replication and repair. Its hub gene, TC010429 “Protein O-mannosyltransferase 1-like Protein” (POMT1), is necessary for pupation, metamorphosis and wing development in *T. castaneum*^35^, and normal muscle formation in *Drosophila*^36^. Finally, the greenyellow module, is implicated in transcription coregulator activity and regulation of gene expression. Its hub gene is TC034166, “Mediator of RNA polymerase II transcription subunit 13-like Protein” (MED13).

**Figure 6.**
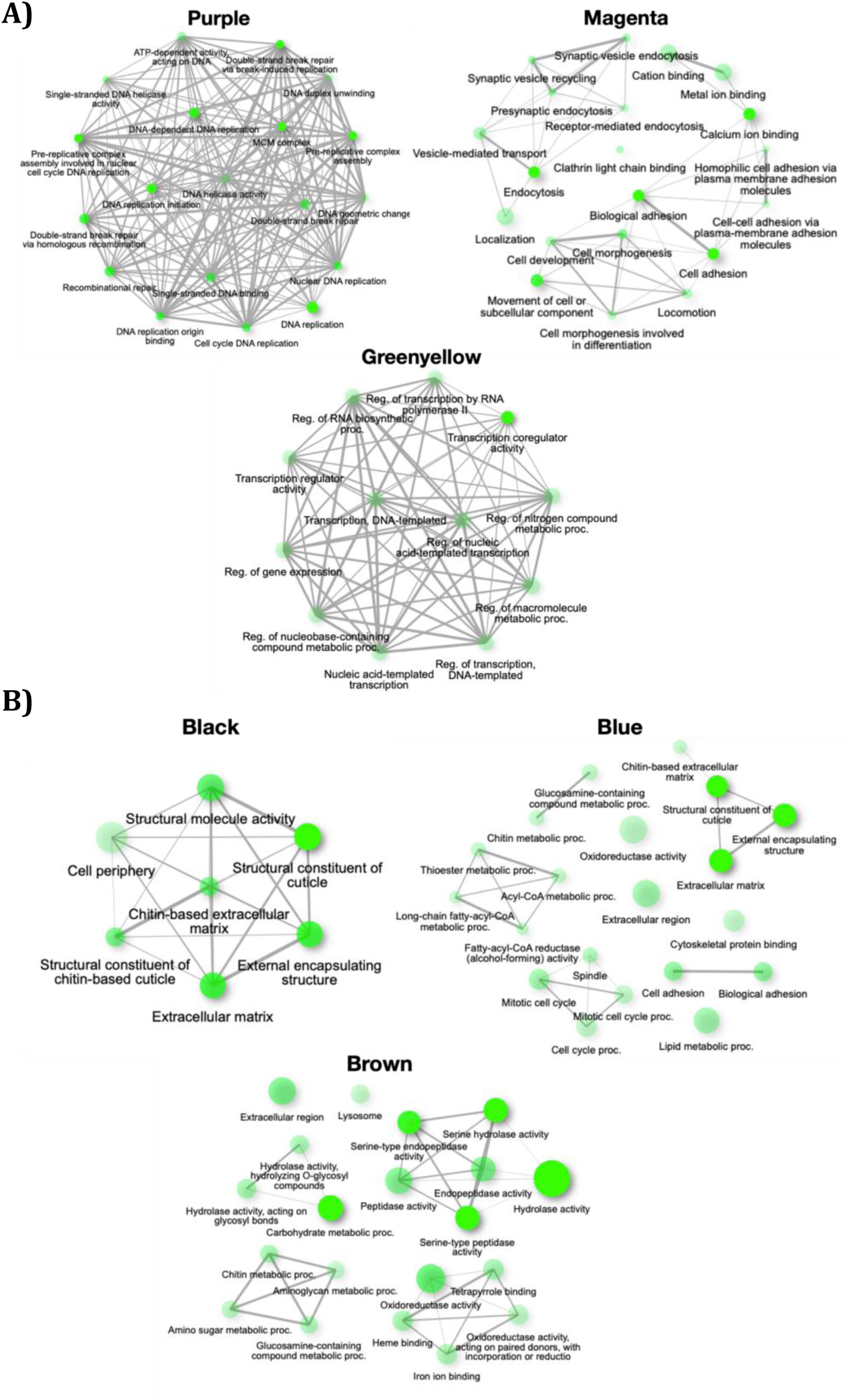
Modules from WGCNA in control samples (without *Btt* exposure). **A)** Networks of enriched GO terms in modules associated with constructed niche treatment and **B)** associated with *Btt* selection. The node size indicates the number of genes. Edges connect related GO terms with their thickness, reflecting the percentage of overlapping genes.

### Divergent constitutive responses to *Btt* selection

In control samples without bacterial exposure, we identified three co-expressing gene modules that were constitutively and differentially expressed between the *pathogen* regimes (Figure 5). The black module (top enrichment term: structural constituent of cuticle), with the hub gene TC013059 (putative defence protein Hdd11) known to be associated with defence against *Bt*^37–39^; and blue module (external encapsulating structure) exhibited a strong negative correlation with *Btt* selection regimes (Figure 6B). While both modules were highly enriched for the structural integrity of the chitin-based larval cuticle, the blue module also included genes involved in the mitotic cell cycle process and metabolism of chitin-related compounds (acyl-CoA fatty acids, lipids, and glucosamine). In contrast, the brown module, which is positively correlated with *Btt* selection, was enriched for hydrolase activity, chitin and carbohydrate metabolic processes, serine-type peptidase activity, oxidoreductase activity, and lysosome (Figure 6B).

### Bacterial exposure induced substantially different gene expression depending on pathogen selection

We also applied WGCNA, to investigate gene expression in response to bacterial exposure across the different regimes. Seven modules showed high correlation with *Btt* selection, of these two were up- and five down-regulated in the *pathogen* selected regimes (Figure S9A). For the upregulated modules, the green module is enriched for serine-type endopeptidase activity, hydrolase activity, and carbohydrate metabolic processes, while the greenyellow module does not contain any significant GO enrichment (Figure S9B). The downregulated modules are enriched for GO terms related to chitin-based extracellular matrix, sugar metabolism, lysosome, peroxisome, and fatty acid metabolisms as well as actin and nucleotide binding, protein modification process, transmembrane ion transport, and gene expression regulation (Figure S9B). No modules were identified to be correlated with different niche regimes or in the pairwise comparison between *niche-free/pathogen* and *niche/pathogen* lines (Figure S9A).

## Discussion

The role of niche construction for evolutionary processes has remained controversial because of the lack of experimental proof. Theory predicts that on the one hand, niche construction might facilitate adaptation, because it creates a feedback loop between genetic and ecological inheritance^5^. On the other hand, a constructed niche might buffer selection pressures, thus causing fixation of deleterious alleles, or stable polymorphisms^5,13,19^. We here designed an evolution experiment to manipulate niche construction, while at the same time selecting for the adaptation to a pathogen. Our results revealed that the extent of niche construction strongly modulated beetle adaptation to its pathogen *Btt*. First, beetles evolving in a constructed niche adapted more rapidly to *Btt*, achieving higher survival rates within just three generations compared to those developing in the absence of niche construction (i.e., without the provision of antimicrobial secretions). This shows that niche construction can facilitate the adaptation to a pathogen in a short time frame. Second, while beetles eventually also adapted to the pathogen in the regime missing niche construction, transcriptomic analysis revealed that resistance evolved via different mechanisms.

Beetles continuously exposed to *Btt* quickly evolved resistance (or tolerance), showing higher post-infection survival, consistent with previous studies on insect exposure to *Bt* or *Bt* products^40–42^. This rapid adaptation could be a case of immediate modification of selection pressures either directly (e.g. antimicrobial quinones) or indirectly (e.g. beetle behaviours or microbiome) by the constructed niche. The constructed niche is rich in antimicrobial stink gland secretions which not only offer protection against harmful pathogens, but might act as signals that prompt immune responses, hence the improved survival rates. The constructed niche might provide a more stable and predictable environment for stronger, more consistent, and directional responses to selection, allowing the earlier development of maximal adaptive potential^43^. An increased immunity to pathogens in response to selection might have plateaued earlier in the constructed niche. Albeit at a slower rate but already after six generations of selection, the beetles not provided with a constructed niche caught up with their counterparts and showed similar levels of survival. The speed of adaptation and fitness varies with resource or environment quality^44,45^. A lesser-constructed niche, as a lower-quality environment, might become an additional stressor to the selection pressure from pathogen exposure, which could slow down adaptation^46,47^.

Against our expectations, we could not observe any immediate trade-off of the evolved survival benefit in either pathogen selection regime (*niche* or *niche-free*). On the contrary, the pathogen-adapted populations underwent earlier pupation than the unselected populations. Developing faster might be a crucial physiological mechanism contributing to the trait of resistance to *Bt* or *Bt* toxins, which is also likely polygenic, as are many life history traits^48,49^. This adaptation might reduce time for feeding on infectious diets, providing an earlier escape from a dangerous environment^50,51^. This difference in developmental speed was especially pronounced in the selection regimes without the provided constructed niche, in which pathogen selected larvae pupated the earliest. A possible explanation for this effect of the lesser-constructed niche is that the additional stressor due to the unstable and unregulated microbiota in the flour intensified the selection pressure for faster development.

In many invertebrates, faster development leads to a reduction in fecundity, because of smaller body size at maturation^52^. Surprisingly, we did not observe lower offspring numbers for the populations that had evolved faster development under pathogen selection pressure. However, fecundity was slightly increased in populations without the constructed niche, which might be a result of being exempted from quinone-mediated chemical interference that inhibits oviposition in constructed niches^53,54^. As shown in other animal species, natal experiences can determine life-history trait expression and future reproductive outputs, potentially acting as a density-dependent feedback mechanism on population growth^55,56^. Females of the *niche-free* populations might perceive a lower population density during their development in the quinone-free non-constructed niche, which would otherwise reduce oviposition to avoid competition for offspring. Although the effect might be relatively small (11% increase in the number of offspring in the *niche-free / pathogen* lines compared to *niche / pathogen* lines), if it persists on an evolutionary time scale, it would quickly accumulate to provide a clear fitness advantage to the populations without niche construction, and might even overshadow the initially faster adaptation in the populations provided a constructed niche.

However, there is a wide range of additional changes in life-history traits (e.g. longevity and late-in-life reproduction) that could trade-off with the evolved resistance and shorter generation time. Furthermore, the cost of developing and maintaining resistance might be separated from other components of host immunity rather than compromising reproduction. In addition, immunity-related trade-offs depend on resource availability^57^; perhaps limited resources (dietary limitation) would give rise to a more substantial trade-off effect in life-history traits for mounting enhanced immunity^58^. Niche construction can amplify or mitigate trade-offs associated with evolved resistance, therefore, we ought to go beyond the larval phenotypic trait, i.e. survival at the larval stage or short-term adult fecundity in this study and explore its consequence under different conditions and timescales.

### Niche construction modulates evolution of resistance mechanisms

Transcriptomic analyses reveal that similar levels of resistance can arise through distinct mechanisms depending on environmental context. In our study, beetles that evolved pathogen resistance without a constructed niche exhibited constitutive changes in gene expression, even in the absence of pathogen exposure, suggesting a form of general, always-active resistance. In contrast, beetles that evolved in the presence of a constructed niche showed inducible expression changes, activated only upon *Btt* exposure. These findings suggest that constitutive responses may offer broader protection in unpredictable microbial environments, where opportunistic pathogens causing secondary infections are more common. But these broader responses likely come with higher metabolic costs due to continuous immune activation^59,60^. Inducible responses, while potentially more limited in scope, are likely more cost-efficient, as they depend on pathogen recognition and are only activated when needed^61^, allowing for a more balanced investment in immunity and other life-history traits. To test these conclusions, future work might investigate the specificity of the evolved lowered susceptibility.

In both the *niche / pathogen* and *niche-free / pathogen* regimes, DEGs before and after *Btt* exposure were enriched for nearly identical Gene Ontology (GO) terms. This suggests that resistance is not driven by entirely distinct mechanisms in the two contexts. Instead, it is likely that similar pathways are involved but are regulated differently, leading to pronounced differences in the number of DEGs within each GO category.

### Constitutive gene expression divergence during resistance evolution

Beetles evolving under *Btt* selection showed a strong downregulation in genes involved in cuticle development, lipid/fatty acid metabolism, mitotic cell cycle process and metabolism of chitin-related compounds, crucial for midgut peritrophic membrane integrity that defends against the invasion of pathogens and gut microbiota into the host haemocoel, aligns with a previous evolution experiment^42^. Downregulation of midgut-specific chitin deacetylase has been reported to decrease susceptibility to baculovirus by reducing peritrophic membrane permeability^62^ and similar downregulation of cuticle genes in various invertebrate species under stress^63,64^, possibly a common adaptive mechanism to abiotic and biotic stressors. Our findings of faster pupation, and elevated expression of serine protease-related and chitin metabolism genes in *Btt*-selected lines highlight the complexity of its regulatory network and suggest the possibility of higher chitin turnover (aided by chymotrypsin-like serine protease) during moulting, which facilitates intestinal maturation and repair. Sustained upregulation of serine protease genes, consistent with findings in *T. castaneum* exposed to live *Bt*^39,65^ and in *Tenebrio molitor* injected with heat-killed *Staphylococcus aureus*^66^, contributes to the activation of innate immune responses, such as hemolymph coagulation, phenoloxidase-mediated melanisation, and antimicrobial peptide (AMP) synthesis via the Toll pathway^67,68^. Constitutive higher expression of serine protease genes in *Btt*-selected beetles might suggest heightened immunity conferred a more efficient or stronger response to infection, prompting the regulatory shift from cuticle-reliant protection to serine protease-related immune responses and faster development.

### Niche-specific transcriptomic responses might aid adaptation to pathogen

Beetles evolving in constructed niches showed constitutively increased expression of genes related to DNA replication, repair, endocytosis, and transcription regulation. This may be an adaptive response to mitigate intense and chronic oxidative stress from benzoquinones, the main component of stink gland secretions^69,70^. The magenta module hub gene, “E3 ubiquitin-protein ligase HUWE1-like protein” (HUWE1) was highly expressed in beetles from constructed niches. It functions in DNA repair and replication stress mitigation^34^, and also aids in foreign protein degradation and immune activation^71^.

Additionally, the induced immune responses to the regular pathogen might also repeatedly expose individuals to excessive oxidative stress from overexpressed immune effectors, such as phenoloxidase and reactive oxygen species (ROS)^72–74^. The adaptation to constructed niches might indirectly select for genes that not only confer enhanced basal resistance to *Btt* but also protection against inflammatory damage from ROS and aid in tissue repair. Alternatively, high oxidative stress from inflammation and constructed niches together might increase the chance of DNA damage and mutation frequency, potentially accelerating the emergence of beneficial mutations that enhance adaptation to other stressors. The combined effects of benzoquinone stress and constant immune activation likely drive adaptive changes in DNA replication and repair pathways, which may explain the high survival rates observed in the *niche / pathogen* regime upon *Btt* exposure.

Genes linked to endocytosis, synaptic vesicle recycling, and biological adhesion were highly expressed in the constructed niche regime, suggesting a role in maintaining cellular homeostasis and facilitating immune and RNAi-related processes^75,76^. These functions are known to be involved in dsRNA uptake in *T. castaneum*^77^ and Toll pathway activation in *Drosophila*^78^. Notably, genes associated with transcriptional regulation, including histone acetyltransferases and mediator complex components, were upregulated in the *niche / pathogen* regime. This aligns with previous findings on epigenetic regulation during transgenerational immune priming^42,65^ and supports the idea that increased histone acetylation may enhance gene expression and contribute to *Bt* resistance^41,79^. Altogether, our results suggest that niche construction can drive heritable changes in immune regulation, potentially accelerating the evolution of host resistance across generations. However, they can also be interpreted as a hint at epigenetic rather than genetic changes underlying the evolved resistance. Thus, future studies should test for epigenetic differences between the evolved populations.

Our findings suggest how niche construction might facilitate adaptation in various ways. First, by sanitising the environment or modulating the surrounding microbiome with stink gland secretions, beetles might be able to reduce infection risk and diffuse selection pressure from pathogens, thereby enhancing their survival and reproductive fitness^25^. The production of these secretions, acting as external immune effectors, could be interpreted as a form of public good^80^ or parental care^25^, leading to a more stable and less stressful environment that allows greater investment into individual immunity^57^. Second, the constructed niche may also contribute to transgenerational immune priming via the inheritance of niches with altered chemical and microbial components, enabling more efficient immune responses against infection. Building on this idea, the constructed niche might systematically reinforce the selection of beneficial traits related to niche construction, which may involve pleiotropic genes that influence not only niche construction, but also host immune defences, as suggested by our transcriptomic findings. Third, this form of niche construction also exposes developing offspring to oxidative stress from benzoquinones. This might either select for adaptive traits that protect individuals from the damage caused by constantly heightened immunity, or it could increase mutations, accelerating the emergence of beneficial traits that enhance resistance to stressors. Future research should explore whether genetic variations differ between populations evolving in constructed and non-constructed niches using a whole-genome resequencing approach. Additionally, detailed chemical and microbial profiling of niches created by selection lines can provide insights into how these major niche components interact and co-evolve with the host. Our findings highlight the active and constructive role of organisms in shaping their environment and offer a mechanistic explanation for rapid evolutionary changes^5,10^. We believe that incorporating niche construction and its reciprocal eco-evo feedback, allows for more precise predictions of evolutionary trajectories as well as a comprehensive framework for understanding the emergence and maintenance of phenotypic variations and speciation. Regarding the ongoing debate about the importance of niche construction^5,10^, we conclude that it plays an important role as it can change evolutionary trajectories, but it remains to be studied in further detail how it scales in comparison to other evolutionary mechanisms, e.g. selection and drift.

For the host, the parasite and their symbionts are part of its biotic environment, while the parasite’s ecological niche is defined by the host–each with their capacity to construct and exert strong selection pressure on one another. In evolutionary medicine and epidemiological studies, the niche construction perspective is crucial for anticipating the outcome of host-parasite coevolution, refining personalized treatments tailored to individual-environment interactions. The concept of niche construction also provides critical insights for conservation efforts under rapidly changing climates or anthropogenic disruptions, as we will have a more realistic view of the environment on population dynamics and fitness. For example, organisms’ capacity to modify or choose an environment might result in enhanced population resilience or limits adaptive potentials to novel stressors.

## Materials and Methods

### Model organisms

#### Insects

*Tribolium castaneum* beetles used here were derived from the CRO1 line^81^, which was originally collected from the wild in Croatia in 2010. The first parental generation consisted of approximately 2,000-3,000 beetles, kept in plastic boxes with heat-sterilised (75 °C) organic wheat flour (Dm, type 550) enriched with 5% brewer’s yeast (hereinafter referred to as “flour”), at 30°C and 70% humidity with a 12h light/dark cycle. The stock population was maintained in non-overlapping generations. The first experimental generation came from a 24h egg-laying period of one-month old adults.

#### Flour conditioning

Upon preliminary characterisation of a pool of candidate genes involved in producing stink gland secretions, we selected the *Drak* gene as the target of the RNAi knockdown to impair proper stink gland development and thereby future niche construction abilities^33^. For each generation, we used siblings of the prior generation as flour conditioning beetles (*niche constructors*, see Figure 1). Pupal knockdown via RNAi (*Drak*-dsRNA 1000 ng/µl dissolved in PBS) impaired the proper development of the stink glands and thus the production of SGS (n*iche-free* regimes). Control beetles (*niche* regimes) received control injections of eGFP dsRNA at the same concentration, which does not impact target gene expression. RNAi treatment in early- to mid-pupal stages followed a previously published protocol^82^ (see supplement for details). One week post injections, one hundred of the enclosed beetles were transferred into jars with fresh flour (0.2g per beetle). The *niche constructors* conditioned the flour for four days, before we sifted it to remove beetles and their eggs, and transferred it into the corresponding jars of the selection lines. Flour exchanges every four days ensured that levels of (volatile) secretions remained constant, as well as sufficient nutrients remaining in the diet. We verified the successful Drak gene knockdown by photometrically measuring secreted quinones^25^ and testing inhibitory abilities of beetle homogenates in a zone of inhibition assay^83^ (see supplements for details).

The evolution experiment consisted of four regimes (Figure 1): Lines undergoing *Btt* selection after developing in constructed niches (*niche / pathogen*), or non-constructed niches (*niche-free / pathogen*), and two control regimes with lines developing in constructed niches (*niche / pathogen-free*) and non-constructed niches (*niche-free / pathogen-free*) without any pathogen selection. Each regime consisted of four replicate lines (with around 100 individuals per line). At the start of the selection experiment, freshly laid eggs were transferred into jars containing the corresponding conditioned flour from *niche constructors* to form the first generation. At day 15 after oviposition, 192 larvae per line were randomly selected and subjected to the *Btt* infection protocol or control treatment. After four days of exposure, 100 surviving larvae were selected and returned to jars with freshly conditioned flour. Exchange of conditioned flour continued until the selection line beetles eclosed as adults and started to produce secretions themselves. After one to two months, these adults were used to produce the next generation of *niche constructors* and *niche inhabitants*.

#### Parasite

*B. thuringiensis morrisoni* var. *tenebrionis* (*Btt*; BGSCID 4AA1) acquired from the *Bacillus* Genetic Stock Centre (BGSC, Ohio State University, USA) was used for the selection and survival assays. The cultivation and oral exposure procedures were modified from the protocol described by Milutinović et al.^81^. Overnight cultures were prepared by inoculating *Btt* from glycerol stock into 3-4 ml *Bt* culture medium containing 15 μl of filter-sterilised salt solution (0.2 μm cellulose nitrate filters, Whatman) and 3.75 μl of 1M CaCl2, and cultured at 30 °C and 180 rpm (see Milutinović et al. 2013 for details on media). The next morning 400 mL *Bt* medium supplemented with 1 mL salt solution and 500 μl of CaCl2 were inoculated with the overnight culture and incubated in the dark at 30 °C and 180 rpm for eight days until completion of sporulation. Another round of the same amount of salt solution and CaCl_2_ was added after four days and for each generation we grew five replicate cultures simultaneously.

Once sporulation had concluded after eight days of incubation, spores were washed and adjusted to a concentration of 2 x 10^10 spores/ml with phosphate-buffered saline (PBS, Calbiochem) and mixed with heat-sterilised flour mixture (0.15 g/mL of spore suspension). The mixture was pipetted (10 μL per well) into flat-bottom 96 well plates (Sarstedt) under sterile conditions and covered with a porous adhesive foil (Kisker Biotech GmbH & Co. KG) and dried overnight at 30°C in the dark. This concentration of spores represents the LD50 (killing 50% of larvae within four days of exposure) for our ancestral population. The diet for the regimes without *Btt* exposure (*niche / pathogen-free* and *niche-free / pathogen-free* treatment groups) was prepared in the same way but without the addition of spores.

### Phenotypic readouts in the F2 generation

Every third generation of ongoing experimental evolution of adaptation to *Btt* in the *niche* and *niche-free* regimes, we performed phenotypic screenings in all replicate lines after two additional generations without niche treatment or *Btt* selection. The relaxation of the selection steps was added to avoid results being compromised by lingering parental effects. We measured the survival post-*Btt* exposure, developmental speed until pupation, and early-life fecundity. All experimental individuals were obtained from two batches of 24 h oviposition from 100 F1 adults in glass jars, as described previously.

#### Survival of *Btt* exposure

To detect the evolution of resistance (or tolerance) against *Btt*, we exposed larvae from all lines of the four selection regimes to 2 x 10^10^ spores/ml of *Btt* spores (LD50 in ancestral population) and recorded their survival for four days. Oral exposure to *Btt* for the survival assay followed the same procedure as in the selection protocol. F2 larvae from each selection line were randomly chosen when 15 days old and 96 individuals each were exposed to *Btt* spores or the control flour only diet.

#### Developmental rate assay

To evaluate whether and to what extent niche construction affects the speed of development, we recorded the pupation date of F2 beetles under *ad libitum* rearing conditions without infection. Fourteen days post oviposition (dpo), 48 larvae from each line were isolated from oviposition jars and individualised into 96-well plates, with each well filled with approximately 0.2 g of loose flour. The larvae were kept individually under standard rearing conditions throughout the experiment. Pupation was recorded daily from 19 dpo to 32 dpo.

#### Early-life, short-term fecundity

To examine the potential cost of different selection regimes for short-term reproductive fitness, we performed a fecundity assay on F2 beetles under *ad libitum* rearing conditions. On the 24^th^ day post-oviposition, most beetles from a single oviposition by F1 beetles were at the pupal stage. We determined their sexes and individualised the F2 pupae. Once all beetles had eclosed and reached sexual maturity, ten single mating pairs of one male and one female from each line were transferred to mating vials containing 5 g of flour for three days. After three days of mating, the adults were removed from the flour. Twelve days after the end of the mating period, living larvae in each vial were counted. The mating assay was performed in two blocks, resulting in 20 single-mating pairs per replicate line.

#### Data analysis of phenotypic data

All statistical analyses and visualisations were performed in R (version 4.2.2^84^) and R studio (version 2022.07.2+576^85^). Generally, we analysed each generation independently and considered selection regimes differing in flour type and bacterial exposure as fixed effects.

For the analysis of the survival of exposure and pupation rate, we used the “Surv( )” function from the “survival” package^86^ (version 2.38) to build the standard survival object and further fitted Cox proportional hazard models after checking that hazard assumptions were met (coxph (survival object ∼ regime)). We tested whether the regimes’ hazard ratio differed significantly from the internal control regime (niche / pathogen-free). Thus, a hazard ratio smaller than one indicates a reduced risk of mortality or a lower pupation rate, respectively compared to the *niche / pathogen-free* control.

To analyse the fecundity data, we used a GLM fitted for a negative binomial distribution with the “lme4” package^87^, with the number of larvae as a response variable and selection treatment and niche type exposed as explanatory variables. We excluded mating pairs with zero offspring, as we could not distinguish between biological responses to stress or treatment and technical issues during the sampling process that might have caused the absence of larvae.

#### Principal component analysis (PCA) on the phenotypic data

Principal component analysis (PCA) was performed to visualise the overall difference between selection populations and explore the relationships between measured phenotypic traits related to fitness, i.e., survival probability (“survival”), day-to-pupation (“development”), number of live larvae produced(“fecundity”). We used the R function “prcomp”, which performed the ordination with standardised means of the mentioned phenotypic traits from each replicate line of selection regimes and calculated the percent of eigenvalues and total variance explained by each principal component (PC). The biplot graphing of the factor loadings and score axes were produced with the “ggbiplot” function from the ggplot2 package^88^.

### Transcriptomic analysis

#### Sample preparation, RNA extraction and RNA sequencing

To understand the potential variations in gene expression resulting from different selection regimes, we performed a transcriptomic analysis of the F2 larvae after nine generations of selection. Samples were prepared in the same way as the survival assay, where the F2 larvae were exposed to control and *Btt* spore (2 × 10^10^cells/mL) diets. Six hours post exposure, we collected live individuals for three replicates of 20 pooled larvae per exposure treatment and selection line (N=96). Total RNA was extracted using a combination of TRIzol (Ambion RNA by Life Technologies GmbH, Darmstadt, Germany), chloroform treatment, and the Total RNA extraction kit with genomic DNA digestion (Promega GmbH, Mannheim, Germany), as described previously^89^. mRNA library preparation (polyA enrichment) and Illumina NovaSeq 6000 PE150 (paired-end 150 bp) were performed by Novogene Co. Ltd. Raw reads containing adaptors with N> 10% and QScores of > 50% bases below 5 were removed. Sequence alignment and mapping were performed using HISAT2 (ver2.0.5) with the reference genome (Tcas5.2.54, http://ftp.ensemblgenomes.org/pub/metazoa/release-54/gtf/tribolium_castaneum/).

#### Differential gene expression analysis

In order to determine significant differences in gene expression between the four selection regimes after nine generations of experimental evolution we made use of the Novomagic software supplied online by Novogene. Before determining differentially expressed genes (DEGs), we calculated Pearson correlations for all replicates of the two treatment groups (6 hrs post *Btt* exposure and control treatment) separately. All R^2^ values of the pairwise comparisons were above 0.8, most above 0.9, confirming that all replicates were technically sound and could be used for further analysis (Figure S9). We ran all further differential gene expression analysis on biological replicates in the Novomagic software (Novogene) using Deseq2^90^. Genes were considered as differentially expressed with any changes in log(2)-fold change and FDR (Benjamini-Hochberg method) adjusted p-value below 0.05. We also performed GO enrichment analysis with Novomagic using GO identifiers from GENEONTOLOGY (https://geneontology.org/).

#### Weighted gene co-expression analysis

Compared with DEGs analysis, which focuses on single genes, WGCNA focuses on a group of co-expressed genes that are involved in various functional pathways. Raw transcriptomic data (GeneID and Gene counts) were split by exposure treatment (control and *Btt* exposure). All genes with counts < 10 in > 75% of the samples were removed, resulting in a reduction in the number of genes in the control samples from 15982 to 11317 and in *Btt*-exposed samples from 16035 to 11403. The counts were transformed using variance stabilisation and outlier samples (CP2_c_2 and CB1_e_3).

Using the entire expression matrix (gene counts), the “cor” function (Pearson’s correlation coefficient) was used to calculate pairwise co-expression values (i.e. the correlation coefficient between any two genes). Genes with similar expression patterns were clustered into ‘modules’ of co-expressed genes which often reflect functionally similar groups of genes. Scale-free, signed, and weighted correlation networks were generated using the R package WGCNA^91^ (v. 1.72-1). A suitable soft threshold of 14 was selected for both the control and *Btt*-exposed samples when the degree of independence was 0.8, using the “pickSoftThreshold” function. The blockwiseModule function was used to construct a signed network using a maxBlockSize of 14000, a “mergeCutHeight” of 0.25 (threshold for merging modules with highly correlated module eigengenes). After module construction, highly correlated modules were merged using dynamic branch cutting with a merging threshold of 0.25.

To identify gene modules that are potentially associated with specific selection regimes, we used the module eigengene values (MEs), the first principal component of the expression matrix for a given module, calculated using the WGCNA algorithm. MEs can be considered the average gene expression levels for all genes in each module. Pearson’s correlation coefficients between the MEs and selection regimes were calculated. The correlation values and p-values are shown in a heat map created using this package. We also identified hub genes in modules of interest. Note that the modules for the non-exposed control samples corresponded to different sets of genes compared to those of the *Btt-exposed* samples.

For gene ontology analysis, genes from the modules of interest were analysed using ShinyGO^92^ (v.0.77) to identify functionally related GO terms and pathways (biological processes, molecular functions, and cellular components). The terms were grouped based on shared genes (kappa score = 0.3). We only included terms with a false discovery rate (FDR) < 0.05, prioritising those with a higher enrichment FDR. We also constructed networks to show the relationships between enriched pathways. Pathways sharing 20% or more genes were linked, with darker nodes indicating more significant enrichment of gene sets, larger nodes representing larger gene sets, and thicker edges signifying a greater gene overlap.

## Data Availability

The sequencing data from this study have been deposited in the NCBI Sequence Read Archive (SRA) under accession number PRJNA1179693.

## Acknowledgements

We thank Zoë Länger, Max Brinkman, Kathrin Brüggemann, and Anke Große-Brinkhaus for help with experiments in the lab. We also acknowledge the intellectual input from Barbara Milutinović and Jaime Anaya Rojas. This research was funded by the German Research Foundation (DFG) as part of the SFB TRR 212 (NC³) – Project numbers 316099922 and 396780003 (to JK)

## Author contributions

L.K.L.: experimental design, data acquisition, data curation, data analysis (Phenotypic data and WGCNA), writing, review and editing; N.S.: experimental design (transcriptome), data analysis (phenotypic data, DEG analysis), writing, reviewing and editing; H.J.: experimental design (ZOI), data analysis (ZOI); J.K.: conceptualisation, experimental design, funding acquisition, methodology, sample collection, project administration, resources, supervision, validation, writing, reviewing and editing. All authors contributed, reviewed and approved the manuscript for publication.

## Competing interests

The authors declare no competing interests.

## Notes

https://www.ncbi.nlm.nih.gov/sra/

### Competing Interest Statement

The authors have declared no competing interest.

