## Supplemental Materials for "Experimental Removal of Niche Construction Alters the Pace and Mechanisms of Resistance Evolution"

### Supplementary Information

#### 1 Results

##### 1.1 Pupal RNAi knockdown caused down-regulation of *Drak* gene expression

We used RT-qPCR to verify the successful knockdown of *Drak* gene expression in adults after being injected with dsRNA during the pupal stage. In preliminary tests, gene expression did not differ between the sexes (data not shown). Across all three analysed time points, the RNAi control beetles injected with *Gfp* dsRNA did not significantly differ in the expression of *Drak* from the untreated naive beetles (Figure S1). However, injection of *Drak* dsRNA significantly reduced target gene expression for all three time points (Figure S1, $F = 21.352$, $df = 3$, $p < 0.001$), demonstrating that knockdown effect lasted for at least 28 days, with no significant difference between the time points.

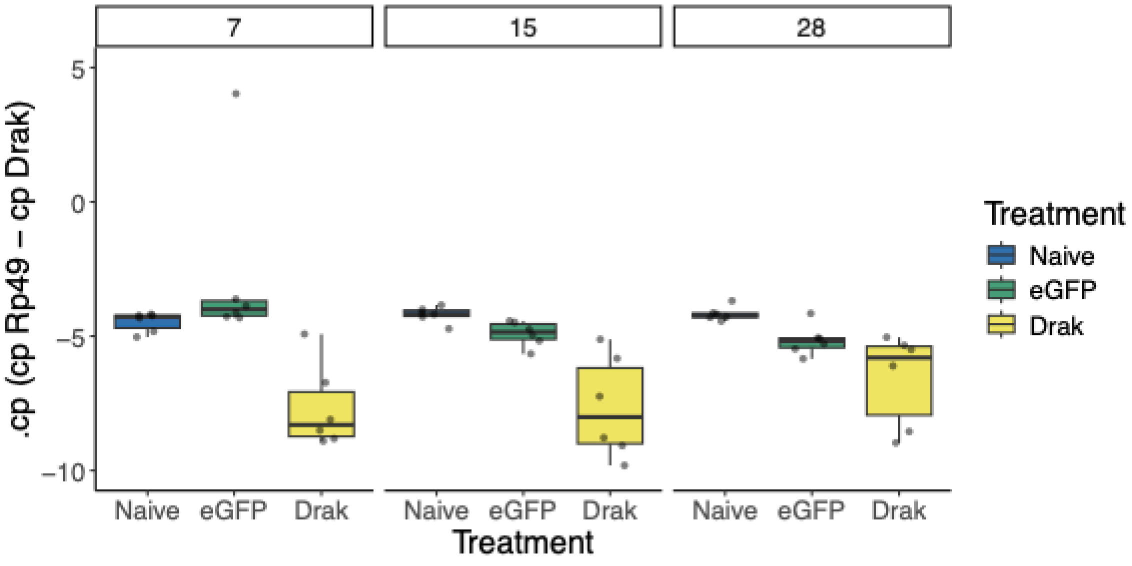


**Figure S1**. Gene expression of *Drak* normalised to the housekeeping gene Rp49 on days 7, 15, and 28 post-injection in *T. castaneum* individuals with control and RNAi treatments (naive, ds*Gfp*-injected RNAi control, ds*Lac2ab*-injected, ds*Drak*-injected). Each box consisted of expression data from six pools (three females and three males) of five beetles per treatment at each time point. Values indicate the difference in the cp-values of the targeted genes and Rp49. The centre lines denote the median, box limits denote the interquartile range, and whiskers extend to the limits. Data points outside the whiskers are outliers. All values were subtracted from those of the blank.

#### 1.2 Knockdown treatment diminished benzoquinones

We measured benzoquinone specific absorbance of collected SGS, to test whether the RNAi treatment reduced secretions in both sexes across all time points. The three-way interaction between time, sex and treatment was not supported, neither were any two-way interactions. There was also no support for a simple effect of the time point on benzoquinone concentration in the samples. However, benzoquinone levels differed by treatment ($\chi^2$ =3,187 = 67.985, df = 3, $p < 0.001$) and sex ($\chi^2$ = 5.208, df = 1, p = 0.022), with males having significantly higher benzoquinone levels than females (t=-2.334, p = 0.0206). As the minimal models showed no interaction between treatment and sex, we further analysed the data separately by sex. In both, females and males, *Drak* knockdown led to a dramatic reduction in benzoquinone levels compared to the control group (Figure S2a; females: t= -9.1914.566, $p < 0.001$; males: t= 4.353, $p < 0.001$). There was no difference in the benzoquinone levels of control and naive beetles (z = 0.381, p = 0.7).

Hydroquinone levels followed the same patterns as benzoquinone levels: there was no support for a three-way interaction between time, sex and treatment, and no support for any two-way interactions, and no support for a simple effect of the time point on hydroquinone concentration in the samples. There was however support for a sex effect ($\chi^2$= 4.91, df = 1, p = 0.024), with males having higher benzoquinone levels than females, as well as an effect of treatment ($\chi^2$ = 111.94100.31, df = 3, $p < 0.001$). In females, the knockdown led to significantly lower benzoquinone levels than in the control group (Figure S2b; Gfp-Drak: t = -9.215.365, $p < 0.001$). Males showed a similar difference (Gfp-Drak: t = 7.111, $p < 0.001$).

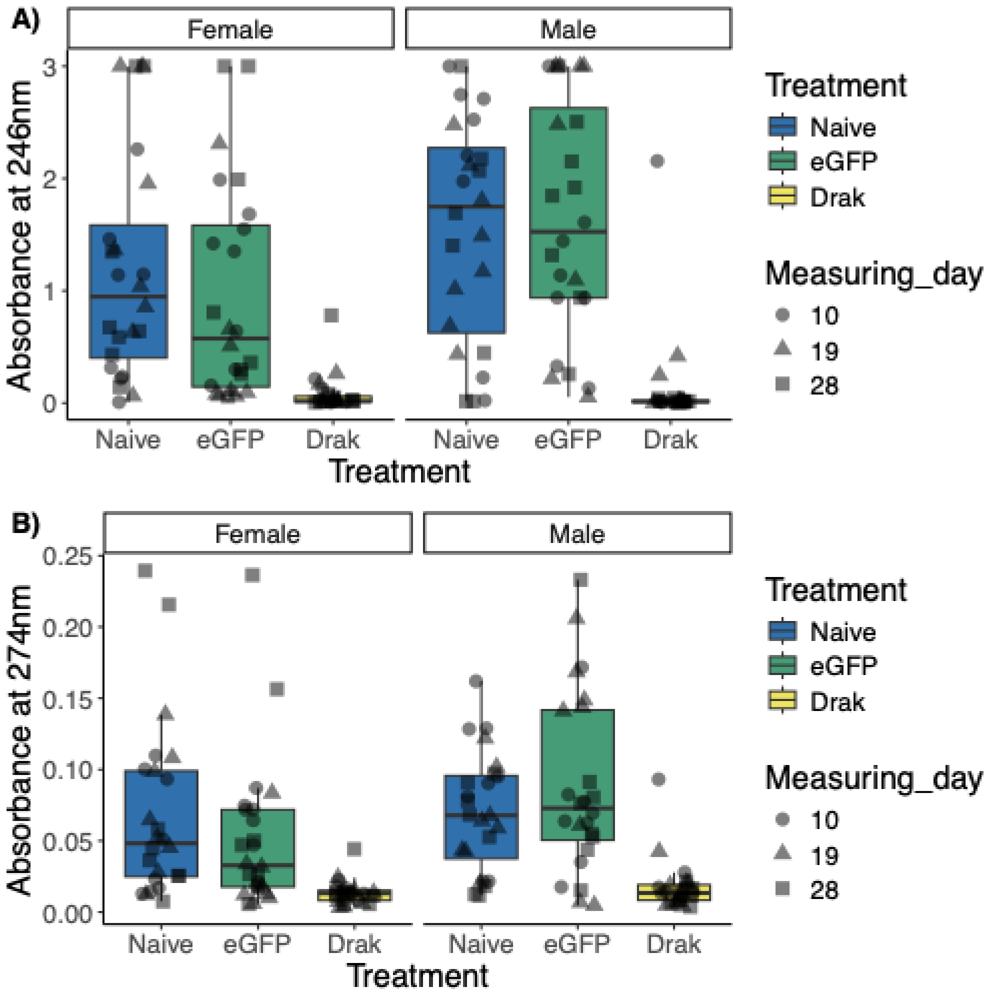


**Figure S2.** Quinone levels in females and males at 10, 19- and 28-days after pupal RNAi. (a) Benzoquinones was quantified by measuring the absorbance at 246 nm while (b) hydroquinone was quantified by measuring the absorbance at 274 nm for each *T. castaneum* individuals, which were either untreated (Naive) or injected with dsRNA (*GFP, Lac2ab, Drak*) during pupal RNAi treatment. The colour of the boxes indicates treatments, and the shape of the data point shows the day post-injection (circle =10; triangle = 19; square = 28). n = 8 per measurement day per sex per treatment. The centre lines denote the median, box limits denote the interquartile range, and whiskers extend to the limits. Data points outside the whiskers are outliers. All values were subtracted from those of the blank.

##### 1.2.1 Knockdown treatment diminished antimicrobial activity

To confirm that the stink gland secretions have antimicrobial properties and that these disappear following a *Drak* gene knockdown, we measured the zones of Btt growth inhibition of beetle homogenates. On average, *Drak* knockdown beetles produced an ZOI with a 0.6 mm radius. This was significantly smaller than and less than a third of the size of the ZOI produced by the control beetles (mean radius: 2.7mm, Wilcoxon rank sum test: W= 242, df=1, $p<0.001$). Additionally, Eleven out of 16 replicates in the knockdown group failed to produce any visible

ZOI, compared to only one replicate in the control group, showing that the knockdown strongly depleted the ability of the beetles to produce antimicrobial secretions.

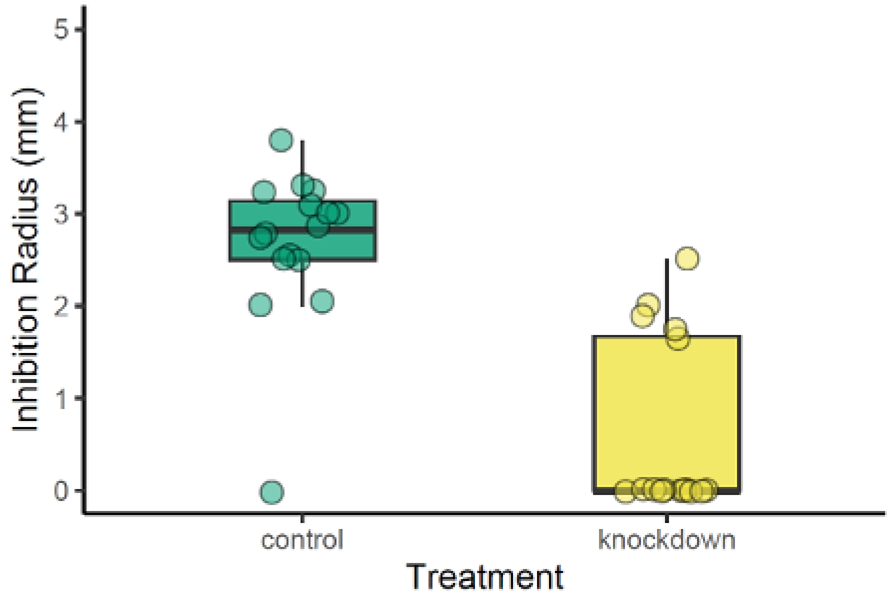


**Figure S3. Zones of *Btt* inhibition produced from beetle homogenates.** Depicted is the visible inhibition radius after 18h of incubation for control beetles (n=16) and beetles that received a pupal *Drak* knockdown (n=16) with the according standard box plot.

#### 1.3 No difference in quinone amounts between selection regimes

To examine the changes in niche-constructing traits upon adaptation, we measured the quinone levels of individual males and females of the F2 offspring of generation 9 from each regime. There was considerable individual variation and no significant effect of niche type (Figure S4; $\chi^2 = 0.31$, df = 1, p = 0.578), pathogen exposure ($\chi^2 = 0.012$, df = 1, p = 0.913), or sex ($\chi^2 = 2.087$, df = 1, p = 0.149). There were no interaction effects between all variables on the mean quinone levels. Although no strong difference was found between the selection regimes, the *niche-free / pathogen* lines had on average a 15% higher amount of quinones than the *niche / pathogen* lines. We observed the largest difference between *niche-free / pathogen* and *niche / pathogen* females, with the former having a 65% higher average amount than the latter (Figure S4).

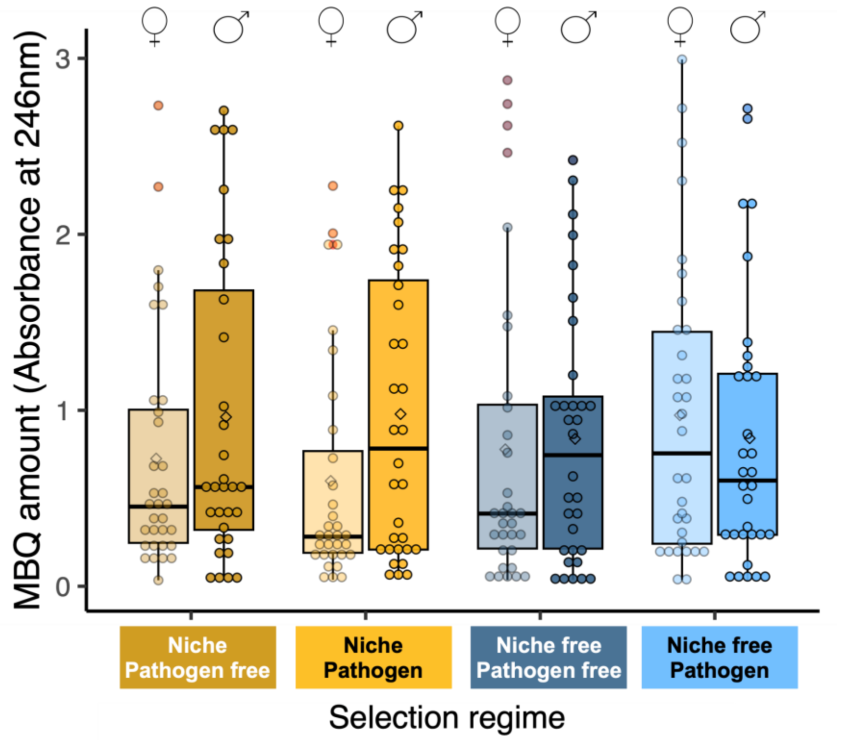


**Figure S4. Benzoquinone levels in male and female adults across all selection regimes.** MBQ levels in the extracts of individual beetles were measured as the peak height at 246 nm. Individual F2 adults from the four replicate lines were grouped together by regime and sex(n = 32). The centre lines of the boxes denote the median, box limits denote the interquartile range, and whiskers show the maxima and minima.

#### 1.4 Differential gene expression analysis

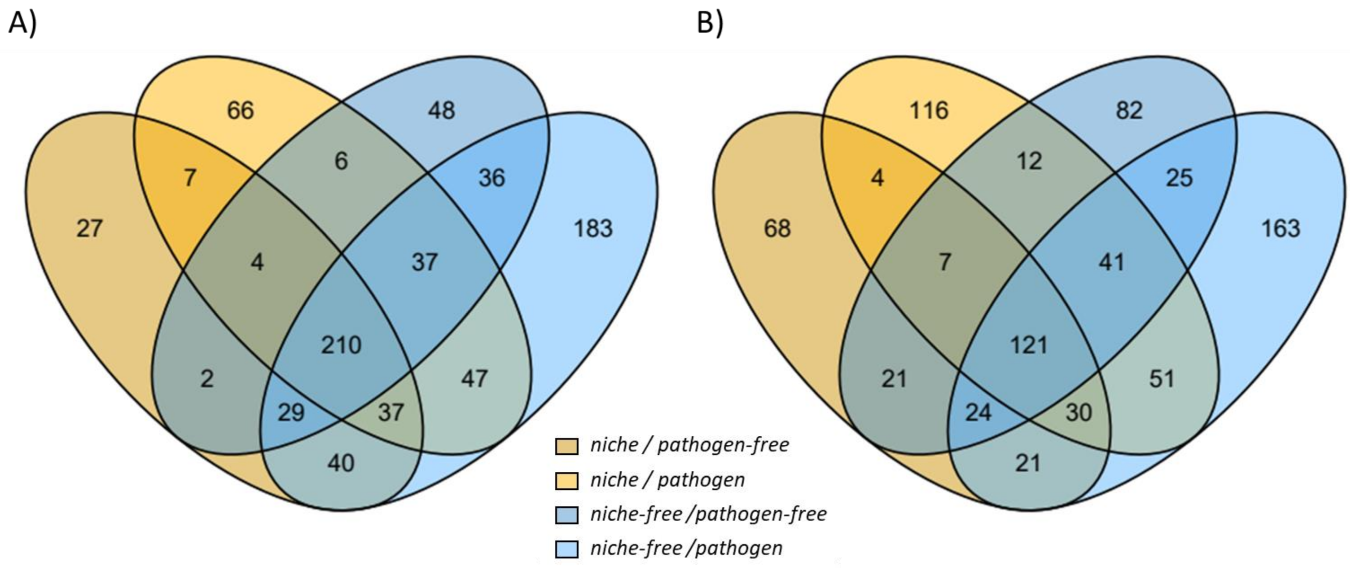


**Figure S5 Venn diagrams of DEGs within the selection regime following *Btt* exposure.** A) significantly downregulated genes B) significantly upregulated genes ($p_{adj}$ <0.05; any log fold change)

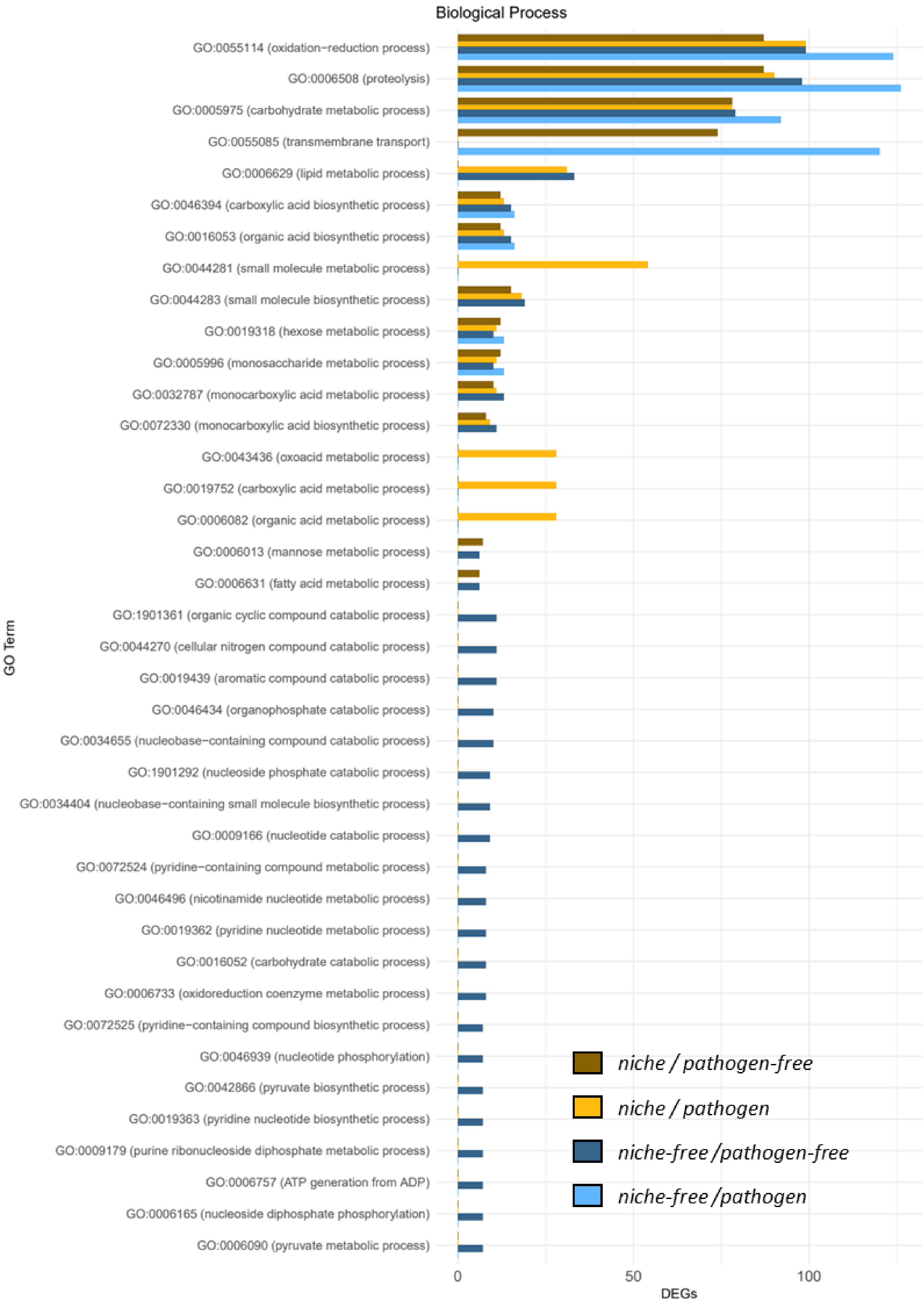
Biological Process
GO:0055114 (oxidation−reduction process)
GO:0006508 (proteolysis)
GO:0005975 (carbohydrate metabolic process)
GO:0055085 (transmembrane transport)
GO:0006629 (lipid metabolic process)
GO:0046394 (carboxylic acid biosynthetic process)
GO:0016053 (organic acid biosynthetic process)
GO:0044281 (small molecule metabolic process)
GO:0044283 (small molecule biosynthetic process)
GO:0019318 (hexose metabolic process)
GO:0005996 (monosaccharide metabolic process)
GO:0032787 (monocarboxylic acid metabolic process)
GO:0072330 (monocarboxylic acid biosynthetic process)
GO:0043436 (oxoacid metabolic process)
GO:0019752 (carboxylic acid metabolic process)
GO:0006082 (organic acid metabolic process)
GO:0006013 (mannose metabolic process)
GO:0006631 (fatty acid metabolic process)
GO:1901361 (organic cyclic compound catabolic process)
GO:0044270 (cellular nitrogen compound catabolic process)
GO:0019439 (aromatic compound catabolic process)
GO:0046434 (organophosphate catabolic process)
GO:0034655 (nucleobase−containing compound catabolic process)
GO:1901292 (nucleoside phosphate catabolic process)
GO:0034404 (nucleobase−containing small molecule biosynthetic process)
GO:0009166 (nucleotide catabolic process)
GO:0072524 (pyridine−containing compound metabolic process)
GO:0046496 (nicotinamide nucleotide metabolic process)
GO:0019362 (pyridine nucleotide metabolic process)
GO:0016052 (carbohydrate catabolic process)
GO:0006733 (oxidoreduction coenzyme metabolic process)
GO:0072525 (pyridine−containing compound biosynthetic process)
GO:0046939 (nucleotide phosphorylation)
GO:0042866 (pyruvate biosynthetic process)
GO:0019363 (pyridine nucleotide biosynthetic process)
GO:0009179 (purine ribonucleoside diphosphate metabolic process)
GO:0006757 (ATP generation from ADP)
GO:0006165 (nucleoside diphosphate phosphorylation)
GO:0006090 (pyruvate metabolic process)
GO Term
0
50
100
DEGs
niche / pathogen-free
niche / pathogen
niche-free /pathogen-free
niche-free /pathogen

**Figure S5 Enriched biological process GO terms following *Btt* exposure** for all four regimes (padj < 0.05; any log fold change). Missing bars indicate no significant enrichment.

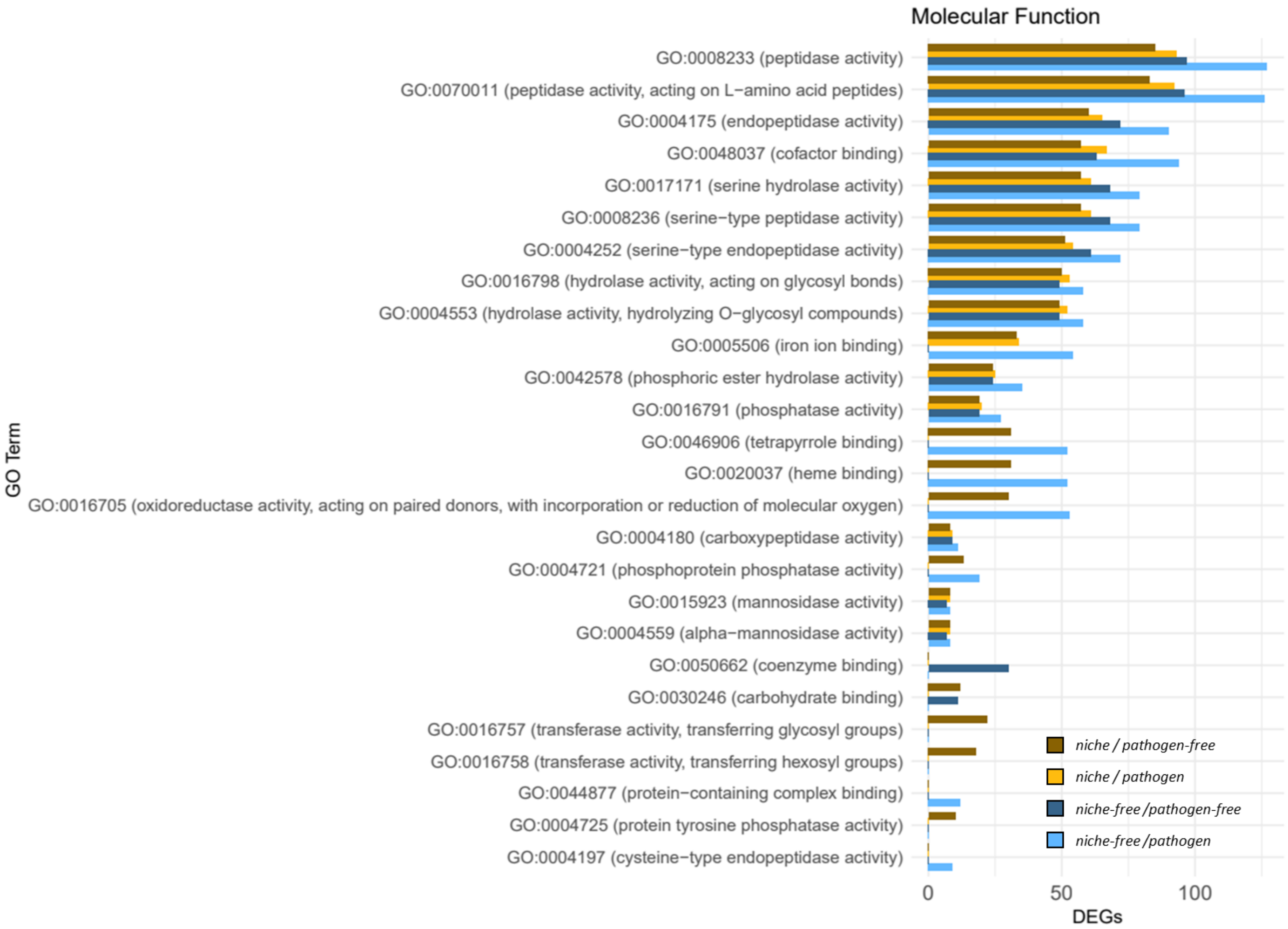


**Figure S6 Enriched molecular function GO terms following *Btt* exposure** for all four regimes (padj < 0.05; any log fold change). Missing bars indicate no significant enrichment.

### WGCNA

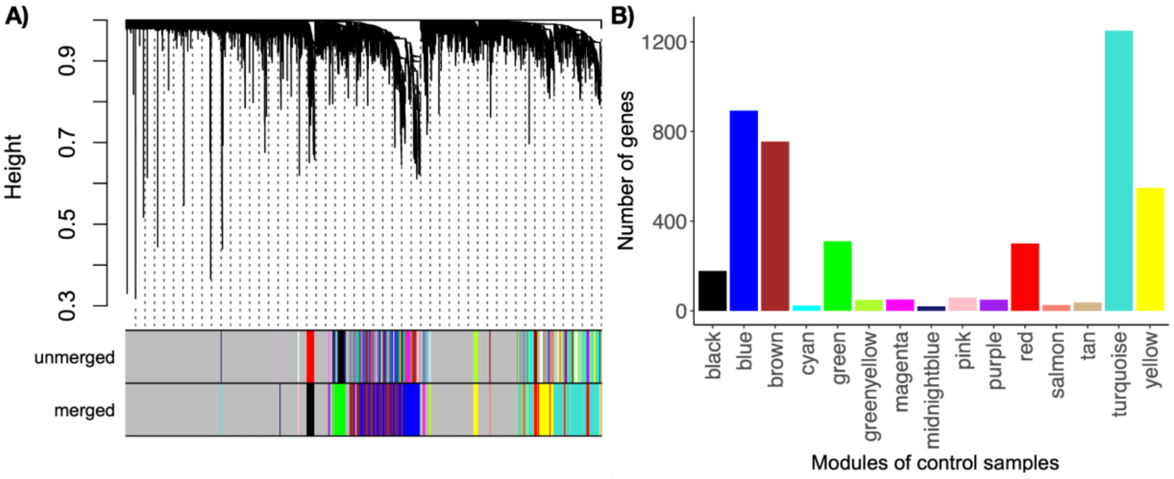

**Figure S7 WGCNA for the control samples. A)** Hierarchical cluster tree of gene co-expression modules. Each branch represents a single gene. The colours underneath represent the modules to which each gene was assigned. **B)** Bar plot of gene counts in each module.

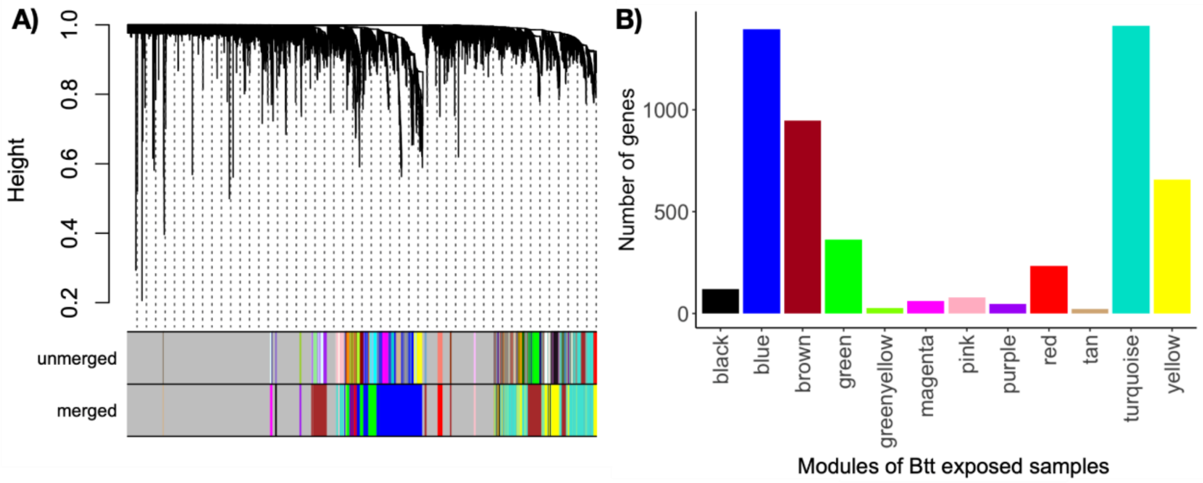


**Figure S8 WGCNA for *Btt*-exposed samples. A)** Hierarchical cluster tree of gene co-expression modules. Each branch represents a single gene. The colours underneath represent the modules to which each gene was assigned. **B)** Bar plot of gene counts in each module.

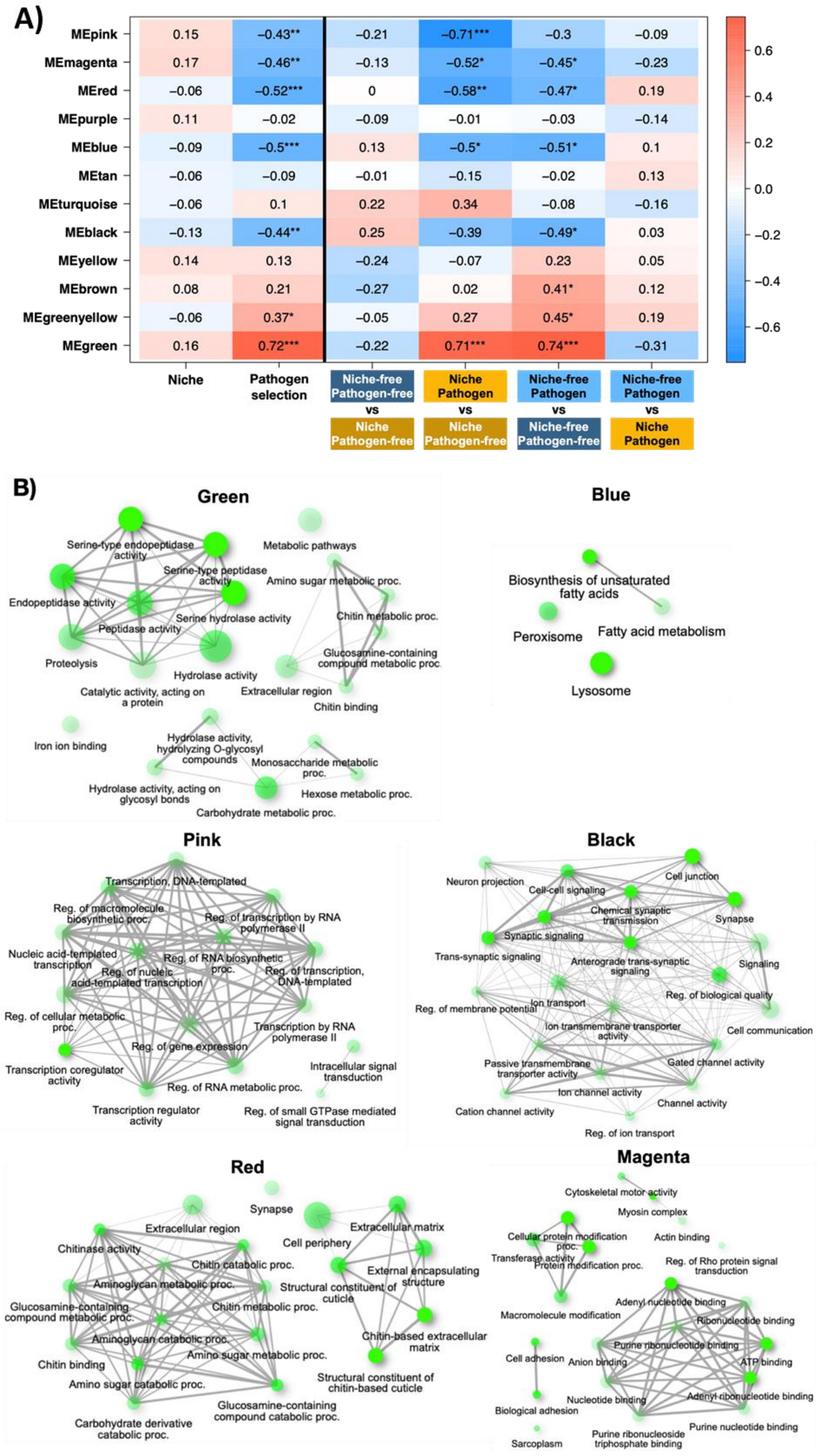
A)
MEpink 0.15 −0.43** −0.21 −0.71*** −0.3 −0.09
MEmagenta 0.17 −0.46** −0.13 −0.52* −0.45* −0.23
MEred −0.06 −0.52*** 0 −0.58** −0.47* 0.19
MEpurple 0.11 −0.02 −0.09 −0.01 −0.03 −0.14
MEblue −0.09 −0.5*** 0.13 −0.5* −0.51* 0.1
MEtan −0.06 −0.09 −0.01 −0.15 −0.02 0.13
MEturquoise −0.06 0.1 0.22 0.34 −0.08 −0.16
MEblack −0.13 −0.44** 0.25 −0.39 −0.49* 0.03
MEyellow 0.14 0.13 −0.24 −0.07 0.23 0.05
MEbrown 0.08 0.21 −0.27 0.02 0.41* 0.12
MEgreenyellow −0.06 0.37* −0.05 0.27 0.45* 0.19
MEgreen 0.16 0.72*** −0.22 0.71*** 0.74*** −0.31
Niche
Pathogen selection
Niche-free Pathogen-free vs Niche Pathogen-free
Niche Pathogen vs Niche Pathogen-free
Niche-free Pathogen vs Niche-free Pathogen-free
Niche-free Pathogen vs Niche Pathogen
0.6
0.4
0.2
0.0
−0.2
−0.4
−0.6
B)
Green
Serine-type endopeptidase activity
Serine-type peptidase activity
Endopeptidase activity
Serine hydrolase activity
Peptidase activity
Proteolysis
Hydrolase activity
Catalytic activity, acting on a protein
Metabolic pathways
Amino sugar metabolic proc.
Chitin metabolic proc.
Glucosamine-containing compound metabolic proc.
Extracellular region
Chitin binding
Iron ion binding
Hydrolase activity, hydrolyzing O-glycosyl compounds
Monosaccharide metabolic proc.
Hydrolase activity, acting on glycosyl bonds
Hexose metabolic proc.
Carbohydrate metabolic proc.
Blue
Biosynthesis of unsaturated fatty acids
Peroxisome
Fatty acid metabolism
Lysosome
Pink
Transcription, DNA-templated
Reg. of macromolecule biosynthetic proc.
Reg. of transcription by RNA polymerase II
Nucleic acid-templated transcription
Reg. of RNA biosynthetic proc.
Reg. of nucleic acid-templated transcription
Reg. of transcription, DNA-templated
Reg. of cellular metabolic proc.
Transcription by RNA polymerase II
Reg. of gene expression
Transcription coregulator activity
Intracellular signal transduction
Reg. of RNA metabolic proc.
Transcription regulator activity
Reg. of small GTPase mediated signal transduction
Black
Neuron projection
Cell junction
Cell-cell signaling
Chemical synaptic transmission
Synapse
Synaptic signaling
Trans-synaptic signaling
Anterograde trans-synaptic signaling
Signaling
Reg. of biological quality
Reg. of membrane potential
Ion transport
Ion transmembrane transporter activity
Cell communication
Passive transmembrane transporter activity
Gated channel activity
Ion channel activity
Channel activity
Cation channel activity
Reg. of ion transport
Red
Synapse
Extracellular region
Cell periphery
Extracellular matrix
Chitinase activity
Chitin catabolic proc.
External encapsulating structure
Aminoglycan metabolic proc.
Structural constituent of cuticle
Glucosamine-containing compound metabolic proc.
Chitin metabolic proc.
Aminoglycan catabolic proc.
Chitin-based extracellular matrix
Amino sugar metabolic proc.
Chitin binding
Amino sugar catabolic proc.
Structural constituent of chitin-based cuticle
Glucosamine-containing compound catabolic proc.
Carbohydrate derivative catabolic proc.
Magenta
Cytoskeletal motor activity
Myosin complex
Cellular protein modification proc.
Actin binding
Transferase activity
Protein modification proc.
Reg. of Rho protein signal transduction
Adenyl nucleotide binding
Macromolecule modification
Ribonucleotide binding
Purine ribonucleotide binding
Cell adhesion
Anion binding
ATP binding
Nucleotide binding
Adenyl ribonucleotide binding
Biological adhesion
Purine nucleotide binding
Purine ribonucleoside triphosphate binding
Sarcoplasm

**Figure S9 WGCNA for *Btt*-exposed samples**. **A)** Heat map of module-selection regime correlations. Cell colours indicate Pearson coefficients, and asterisks denote significance levels. **B)** Networks of enriched GO terms in modules associated with *Btt* selection. The node size indicates the number of genes. Edges connect related GO terms with their thickness, reflecting the percentage of overlapping genes.

### 2 Methods

#### 2.1 RNAi knockdown

DsRNA of *Drak* (TC031247) was manufactured by Eupheria and dissolved in PBS with a concentration of 1000 ng/µl. DsRNA of the enhanced Green Fluorescent Protein (eGFP) served as a procedural control in our experiments (concentration adjusted to (1000 ng/µl, in PBS). EGFP-dsRNA was generated in our lab according to existing protocol (Posnien et al., 2009) and stored at -20 ºC until use.

**Table 1** dsRNA sequences for the genes of interest

| Gene | Sequence |
|---|---|
| *Drak* (TC031247) | TTTGGACTTAAAACCGCAGAATTTACTGCTCTCGATCGAGGATAAC<br>TGTGATGATATCAAGCTCTGTGATTTCGGCATATCGAAGGTTTTGG<br>AGCCCGGCGTTAAAGTACGGGAGATTATAGGTACTGTAGATTATG<br>TAGCGCCGGAAGTGCTTAGTTACGACCCGATCTGTCTGTCGACTGA<br>TATTTGGTCGATTGGAGTTTTGGCGTATGTGCTTTTGTCGGGCTATA<br>CGCCGTTTGGGGCCGACGATAAGCAACAGACTTTTCTTAATATATC<br>AAAGTGTGCCTTATCTTTCGAGCCAGACCATTTCGAGGATGTTTCC<br>TCGCCGGCTATAGACTTTATCAAGAGTGCCTTGGTCACGGATCCCA<br>GGAAGCGTCCGACCGTCCACGAGCTGTTGGAGCACCCGTGGATTT<br>CGCTGAAGTGCCCCCTCCTGCCGGCGCTGCCGCTGAAGCCTTCGGA<br>GTCCGTTCCGTCGATAAA |
| eGFP | TGGAATGAGCATCTGATGCTGCTCTATCGGCGCCTTGTGATGGATG<br>CCAGGGCGATGCCACCTACGGCAAGCTGACCCTGAAGTTCATCTG<br>CACCACCGGCAGCTGCCCGTGCCCTGGCCCACCCTCGTGACCACCC<br>TGACCTATGGCGTGCACTGCTTCAACCACTACCCCCACCACATGAA<br>GCCCTTGACTTCTTCAAGTCCTCCATGCCCGAAGGCTACTTCCGGG<br>AGCGCACCATCTTCTTCAAGGACGACGGCCACTACAAGACCCGCG<br>CCGAGGTGAAGTTCGAGGGCGACACCCTGGTGAACCGATCTGCAG<br>AATTCCAGCACACTGGCGGCCGTTACTAGTGGATCCCCAGCTCGGT<br>ACCAAGCTGGATGCATAGCTTGAGTATTCTATAGTGTCACCTAAAT<br>CCTATAGTGAGTCCTATTACAATTCAA |

Pupae in the early- and mid-pupal stages were immobilised on glass slides (diagonal, 78 × 26 mm) with glue (Fixogum, Marabu GmbH) before receiving approximately 0.5 µl  of dsRNA of either the *Drak* gene or the eGFP gene as a procedural control. A batch of beetles did not receive any injection ("naive"), but was similarly handled. The dsRNA was injected dorsolaterally (at the ventro-lateral position between the 3rd and 4th abdominal segments) into each of the insects using pulled glass capillaries (borosilicate glass 3.3, ends cut, 100 mm length, 1 mm outside diameter, 0.58 mm inside diameter, Hilgenberg) and a micro-injector (FemtoJet, Eppendorf AG, with settings of pi = 200 hPA and pc = 19 hPa) under a dissecting stereomicroscope. After injection, all pupae were then kept facing-down in petri dishes with approximately 10 g of flour and incubated under standard rearing conditions. We checked daily for newly eclosed adults and individualised them in 96-well plates to prevent mating and injuries until the further experiments.

#### 2.2 Verification of knockdown efficiency

We assessed the magnitude and duration of the knockdown effect on days 7, 15, and 28 post injection. For this, six pools of three adult beetles from each sex and treatment group were collected into 2 ml safe-lock Eppendorf tubes and frozen in liquid nitrogen. Afterwards, the samples were stored at -80 °C until RNA extraction, cDNA reverse transcription, and RT-qPCR, as described previously (Schulz et al. 2022). We also verified the knockdown effect using a non-overlapping secondary construct (data not shown).

To further confirm the successful inhibition of the release of SGS after *Drak* gene knockdown, we measured collected secretions from treated beetles as described in Joop et al. (2014) and photometrically measured their quinone specific absorbance at the wavelengths, characteristic for the different quinones. For each selection line, eight female and eight male beetles (36-day-old virgin adults) were randomly selected and individually transferred to 1.5 ml safe-lock reaction tubes (Eppendorf AG). They were then placed on ice for five minutes to prompt relaxation of the stink gland reservoir's closing muscles and the release of all secretions from the glands (Loconti & Roth, 1953; Unruh et al., 1997). We then euthanised the beetles by freezing them at -20 °C for 10 min. 160 µl acetonitrile (ROTISOLV ≥99,98 % Ultra LC-MS, Carl Roth) was pipetted into each tube and subsequently incubated overnight at 4-7 °C in the dark. The next day, tubes containing the extracts were shaken in a thermomixer (5 min at 5 °C at 1000 rpm) to ensure the complete dissolution of quinones and to wash residual quinones off the inner walls of the tubes. To spin down all beetle body parts and flour, the samples were centrifuged at 10,000 rpm at 5 °C for 5 min. Next, 120 µl of the extract (supernatant) was pipetted into a pre-cooled Quartz-96-well microliter plate (Hellma, Germany) and quinone concentration was measured with a Tecan plate reader (Tecan Deutschland GmbH, Tecan infinite 200). A control of acetonitrile undergoing the same extraction process, and a blank control containing the solvent (acetonitrile), were pipetted into the wells and measured along with the samples. To detect the level of benzoquinones, the absorbance at 246 nm, i.e., the optical density at which benzoquinone peaks, as well as that of a spectrum ranging from 200 nm to 450 nm, was measured to ensure the absence of contaminants. Gene expression data were processed with the REST2009 software, as described previously (Eggert et al. 2014). Quinone data were analysed with generalised linear models fitted for a gamma distribution (log-link), with absorbance at 246 nm and 274 nm as response variables, and treatment and sex as fixed factors (best model excluded measurement day).

Additionally, we performed a zone of inhibition (ZOI) assay, to measure the ability of wash-offs from the different *niche constructors* in inhibiting the growth of *Btt* and to test whether the ZOI is reduced following RNAi knockdown. For this assay, we followed the protocol from Roth *et al.* (2010). In short, *Btt* from frozen glycerol aliquots (-80 °C) were incubated on standard LB agar overnight at 30°C. For each of two replicates, 5 ml sterile LB medium were inoculated with a sample from a single colony and incubated overnight at 180-200 rpm and 30 °C. For the preparation of bacteria plates for the assay, we used 6ml LB medium with 1% agarose and added spores for a final density of 4 x $10^5$ cells/ml. Uniform growth was exemplary checked by incubating one plate overnight.

For the assay, two-month-old adult *niche constructor* beetles from all four selection regimes (after receiving the respective RNAi treatment during the pupal stage) were individually frozen and stored at -80 °C until the experiment. To create beetle homogenates 30 µl insect Ringer solution supplemented with phenylthioureic acid (PTU, to inhibit melanization) was added to frozen beetles. All samples were homogenized using metal beads at 30 Hz/s for 2 minutes in a ball mill (Retsch MM301, Retsch, Germany). They were subsequently centrifuged at 6500 rpm and 4 °C for 10 min and the supernatant was used in the zone of inhibition assay.

Before adding the samples, eight holes with a diameter of 2 mm were punched into the bacteria agar plates using autoclaved 5 ml pipette tips. Into these holes, 2 µl of supernatant was added. Each plate included two positive controls (25 µg/ml and 50 µg/ml streptomycin) and one negative control (Ringer + PTU). The samples were pseudorandomly distributed across the wells with no single sample type appearing twice on one plate. Following pipetting, the plates were inverted and incubated at 30 °C for 18 hours. The zones of inhibition were measured using a manual caliper by measuring the radius two times and taking the mean. Additionally, agar height was measured for each plate. For the statistical analysis, we used a Wilcoxon rank sum test to assess differences between the secretions of knockdown and control beetles, because the residuals for the measured radii did not follow a normal error distribution.

#### 2.3 Statistical analysis

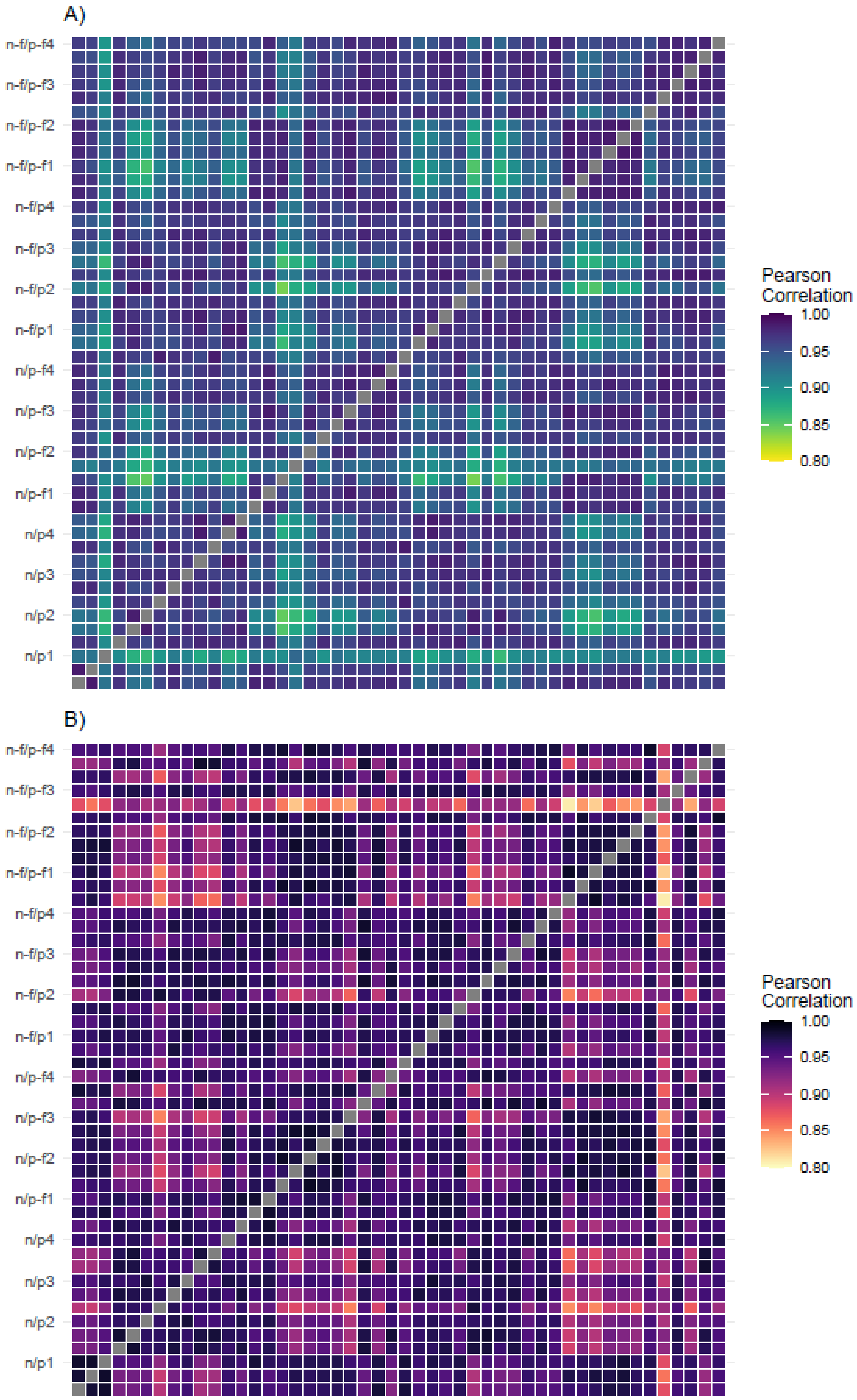

**Figure S9 Pearson correlations between all RNAseq samples**. **A)** without and **B)** with *Btt* exposure. Colors are depicting $R^2$ values between 0.8 and 1.
